# Single-cell data reveal heterogeneity of resource allocation across a bacterial population

**DOI:** 10.1101/2024.04.26.591328

**Authors:** Antrea Pavlou, Eugenio Cinquemani, Corinne Pinel, Nils Giordano, Mathilde Van Melle-Gateau, Irina Mihalcescu, Johannes Geiselmann, Hidde de Jong

## Abstract

Ribosomes are responsible for the synthesis of proteins, the major component of cellular biomass. Classical experiments have established a linear relationship between the fraction of resources invested in ribosomal proteins and the rate of balanced growth of a microbial population. We extended the study of ribosomal resource allocation from populations to single cells, using a combination of time-lapse fluorescence microscopy and statistical inference. We found a large variability of ribosome concentrations and growth rates in conditions of balanced growth of the model bacterium Escherichia coli. Moreover, the ribosome concentrations and growth rates of individual cells are uncorrelated, contrary to what would be expected from the population-level growth law. A similar large heterogeneity was found during the transition of the bacteria from a poor to a rich growth medium. Whereas some cells immediately adapt ribosomal resource allocation to the new environment, others do so only gradually. Our results thus reveal distinct strategies for investing resources in the molecular machines at the heart of cellular self-replication. This raises the interesting question whether the observed variability is an intrinsic consequence of the stochastic nature of the underlying biochemical processes or whether it improves the fitness of Escherichia coli in its natural environment.

## Introduction

Microbial growth involves the conversion of nutrients from the environment into biomass. The main component of biomass are proteins, which also play a major role in the synthesis of new biomass by functioning as enzymes in metabolism and by constituting the molecular machinery responsible for the synthesis of proteins and other macromolecules. Microbial growth therefore requires the coordinated investment of cellular resources in different categories of proteins. The basic principles of this coordination have been described by so-called growth laws [1-3]. Growth laws relate the growth rate of the cells to the abundance of different categories of proteins. In other words, they quantitatively describe the resource allocation strategies adopted by microbial cells.

A classical growth law states that there exists a linear relationship between growth rate and the mass fraction or concentration of ribosomal proteins [4-11]. Ribosomes are probably the most important protein category for two reasons. First, they are responsible for the synthesis of all proteins in the cell. Second, they are themselves very costly to make: ribosomes constitute up to 40-50% of the total protein mass in *Escherichia coli* [12, 13]. The ribosomal growth law has been shown to hold over a large range of growth rates and for a wide variety of microorganisms, and has been generalized to cases of reduced translation rates by antibiotic treatment [9] and different temperatures [14]. Simple mathematical models explain the growth law from basic hypotheses on cellular resource allocation under the assumption that cells have evolved to maximize their growth rate [9, 11, 14-21].

The ribosomal growth law applies to populations of microorganisms in a state of balanced growth. In natural environments, however, growth is rarely balanced and microorganisms have to cope with regular changes in nutrient availability and other perturbations [22]. Theoretical models have predicted how microorganisms dynamically adapt the synthesis of ribosomes during shifts between balanced growth on a poor or a rich carbon source [18, 23, 24]. A recent proteomics study reported changes in the (relative) abundance of ribosomes and other proteins during such upshifts and downshifts [23]. This has motivated generalizations of the growth law to unbalanced scenarios, relating ribosomal protein fractions to observed jumps in growth rate after a nutrient upshift [25] (see also [26]).

The quantification of resource allocation on the single-cell level has not been attempted thus far. It is well-known, however, that individual cells within a microbial population do not behave identically, but display variable phenotypes [27-33]. How do the resources allocated to the synthesis of ribosomal proteins, both during balanced and unbalanced growth, vary over the individual cells of an isogenic population? In order to answer this question, we constructed chromosomal reporter systems for monitoring ribosome expression in the model organism *Es-cherichia coli*. We measured the ribosome concentration in individual cells growing on a rich or a poor carbon source, as well as changes in ribosome concentration during upshifts and downshifts between these carbon sources, over extended periods of time (>80 generations). Moreover, we developed a method for the statistical inference of dynamic strategies for ribosomal resource allocation from the single-cell, time-course data thus acquired.

We found that, during balanced growth, the bacteria display a wide variety of ribosome concentrations that are mostly uncorrelated with the single-cell growth rate. This would not be expected if bacteria had optimized costly ribosome expression to precisely match the protein synthesis rate required for a certain growth rate. It would be consistent, however, with the existence of a variable ribosome reserve which the bacterial cells may exploit to speed up adaptation to sudden changes in the environment [25, 26, 34]. During the upshift from a poor to a rich carbon source, we indeed observed that cells with a higher pre-shift ribosome concentration more rapidly adapt resource allocation to the new environment. However, fast adaptation of resource allocation as compared to slow adaptation does not translate into a higher growth rate during the upshift. These results do not suggest that fast adaptation of ribosomal resource allocation brings a fitness advantage in the growth transition considered here, but the observed variability may be advantageous in other situations that *E. coli* confronts in its natural habitat.

In this study, we thus quantified for the first time, using a combination of reporter genes and powerful statistical inference algorithms, bacterial resource allocation strategies on the single-cell level. The results reveal a surprising variability in the allocation of resources to ribosomes, the most costly molecular machine in bacterial cells, during both balanced and unbalanced growth. This raises fundamental questions on the role of variable resource allocation strategies in shaping the growth of a bacterial population and its adaptation to changing environments. Given the importance of growth and adaptation in biomedical and biotechnological applications, we expect our findings to have practical implications as well.

## Results

### Single-cell experiments to monitor changes in ribosome concentrations

In order to quantify the ribosome abundance in single *E. coli* cells, we tagged one of the ribosomal subunits with a green fluorescent protein (GFP). In particular, following previous work [35], we constructed on the chromosome of the *E. coli* strain BW25113 [36] a translational fusion of the *rpsB* gene, coding for the S2 subunit, and the gene encoding the fast-folding GFPmut2 (*Materials and methods*). The resulting strain, expressing the fusion protein RpsB-GFPmut2, was called Rib (*Materials and methods*).

We verified for the reporter strain, in batch experiments using a variety of different media, that the growth rates of the population are not significantly different from the corresponding growth rates of the wild-type strain (Fig. S1). Furthermore, we quantified in these experiments the fluorescence emitted by the cultures during balanced growth and normalized the fluorescence by the optical density, yielding a quantity proportional to the ribosome concentration on the population level. When plotting this quantity as a function of the growth rate, for the different conditions, we found a linear relationship in agreement with the growth law for ribosomes (Fig. S2). These control experiments demonstrate that the constructed reporter strain provides a reliable read-out of ribosome concentrations over a broad range of growth conditions.

In order to quantify ribosome concentrations in single cells, we used an experimental setup in which cells of the Rib reporter strain are grown in a mother machine and monitored using time-lapse fluorescence microscopy (*Materials and methods*). Mother machines are microfluidics devices that allow cells trapped at the bottom of dead-end microchannels, so-called mother cells, to be followed over hundreds of generations in well-controlled conditions of growth [37]. We submitted cells in the mother machine to upshifts from a poor growth medium (minimal medium with acetate) to a rich growth medium (minimal medium with glucose). After each upshift, the bacteria were left to grow for a sufficient number of generations (> 10) to reach a state of balanced growth in the new medium. Each experiment consisted of at least two upshifts, separated by corresponding downshifts from glucose to acetate (Fig. S3). During these experiments, we acquired phase contrast and green fluorescence images at regular time-points (*Materials and methods*). Using existing image analysis software [38], we segmented the mother cells and recorded for each cell at each measurement time the cell length and (average) fluorescence intensity (Fig. S4).

This experimental setup allowed us to monitor the accumulation of ribosomes in single cells over long periods of time (several days) with high sampling density (measurement every 11-13 min). As illustrated in Fig. 1, the experiments reproduce a number of well-known features of *E. coli* growth. After the upshift from acetate to glucose, the cells are seen to divide faster, corresponding to the higher growth rate on glucose, the preferred carbon source of *E. coli*, and have a larger newborn cell length on average [32]. The fluorescence intensities increase after the upshift and stabilize at a higher level, reflecting that the higher growth rate on glucose comes with a higher ribosome concentration [6, 8, 9]. Following the downshift, a lag phase is observed before the cells resume growth on acetate [39]. These observations are reproducible in independent replicate experiments (Fig. S5).

**Figure 1:**
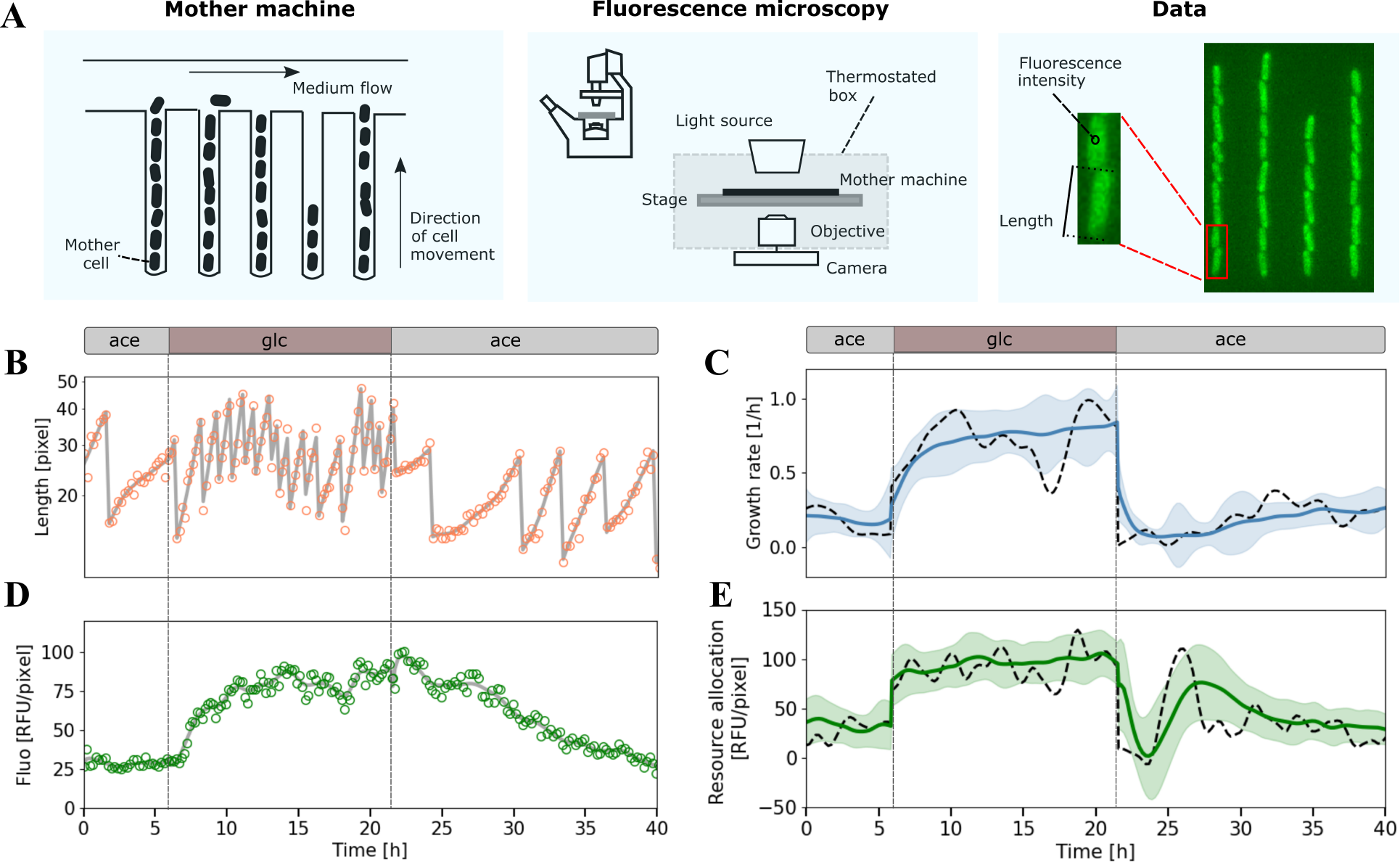
Measured and estimated quantities in mother machine experiments with a ribosomal reporter strain. **A**. Schematic outline of the mother machine and the use of fluorescence microscopy to quantify cell length and fluorescence intensity over time. The fluorescence intensities of a cell reported in the text are the means of the intensities of the pixels of the cell. **B-C**. Cell length in log scale (orange dots) and green fluorescence intensity (green dots) of individual bacteria of the Rib strain, carrying a fusion of the ribosomal subunit S2 and the green fluorescent protein GFPmut2. The experiment consisted in several consecutive upshifts and downshifts (vertical dashed lines) between minimal media with glucose (glc) or acetate (ace). **D-E**. Cell length and fluorescence intensity measurements were used to estimate growth rates and resource allocation strategies, respectively, using appropriate statistical inference methods (Materials and methods). The gray solid curves in panels B and D represent the fits of the single-cell data obtained from the inference methods. The black dashed curves in panels C and E represent the corresponding estimates of the growth rate *µ*(*t*) and the resource allocation strategy *α*(*t*)*/β* for this same mother cell. Blue and green solid curves represent the mean of the estimates over all cells considered in the experiment and confidence intervals are given as two times the standard deviation.

### Inference of dynamic resource allocation strategies

We exploited the data obtained from the mother machine experiments to quantify (i) resources allocated to the production of ribosomes and (ii) changes in resource allocation after a nutrient upshift or downshift. To this end, we first formulated a simple model describing the dynamics of the ribosome concentration in a mother cell. The model is a variant of previously published models ([16, 18] and Text S1). The model assumes that the ribosome concentration evolves continuously across cell divisions, which is a reasonable approximation given the high number of ribosomes in *E. coli* cells growing on glucose or acetate (between 8000 and 15000 8, 12). The dynamics of the ribosome concentration *r* as a function of time *t* is defined in terms of the time-varying growth rate *µ*(*t*), the protein degradation rate *γ*, the total protein concentration 1*/β*, and the resource allocation strategy *α*(*t*). The total protein concentration is assumed constant, which is a good approximation for the conditions considered here [40] :

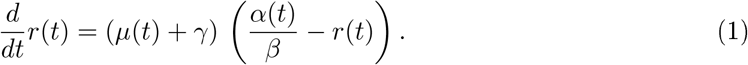

The model expresses that the instantaneous change in ribosome concentration arises from the difference between the rate of decay of ribosomes due to growth dilution and degradation ((*µ*(*t*)+ *γ*) *r*(*t*)) and the synthesis rate of new ribosomes as a fraction *α*(*t*) of the total protein synthesis rate (*µ*(*t*) + *γ*)*/β*. In other words, resource allocation corresponds to the distribution of the total protein synthesis rate, the capacity to make new proteins, over different protein categories, and *α*(*t*) denotes the fraction allocated to ribosomes.

How do the data provided by the mother machine experiments, that is, the time-lapse measurements of the length and (average) fluorescence intensities of a cell, relate to the above model? In brief, the cell length measurements allow us to infer an estimate of the time-varying growth rate *µ*(*t*), in units 1/h, while the fluorescence intensities report on the time-varying ribosome concentration, expressed in relative fluorescence units per pixel (RFU/pixel). Together with the measured degradation constant *γ* of the fusion protein, obtained in previous work [41], this information allows us to infer an estimate of *α*(*t*)*/β*, in units RFU/pixel. While the absolute value of this ratio is difficult to interpret, because the value of 1*/β* is not precisely known for our strain, the changes in *α*(*t*)*/β* following a nutrient upshift or downshift will inform us about relative changes in *α*(*t*).

In order to estimate single-cell growth rates from cell length data, we developed a dedicated method based on work from statistical signal processing (Fig. 1A, *Materials and methods*, and Text S2). Contrary to many estimation methods applied to microfluidics data, we do not assume that growth rates are constant between two cell divisions. This is critical for monitoring medium shifts, when bacteria change their growth rate dramatically within one generation (Fig. S6). The method uses regularization to cope with measurement noise in the data, thus penalizing rapid changes in growth rate. The regularization parameter was determined from the data by means of cross-validation. Fig. 1B and D illustrate the estimation of the growth rate of a typical mother cell across an upshift and a downshift, and the corresponding fit to the length data. The mean growth rate over all mother cells considered in the experiment increases from 0.27 *±* 0.02 h^*−*1^ on acetate to 0.79 *±* 0.03 h^*−*1^ on glucose, in agreement with values measured in batch for the BW25113 wild-type strain [42, 43].

For the estimation of single-cell resource allocation strategies, we formulated the inference of *α*(*t*)*/β* from the growth-rate estimations and the fluorescence measurements as a Bayesian estimation problem and solved this problem by means of Kalman smoothing [41, 44]. The overall information flow of the procedure is summarized in Fig. 2, while more details can be found in the *Materials and methods* and Text S2. The method uses a variant of the model of Eq. 1 that accounts for maturation of the fluorescent reporter. Ignoring maturation may distort the inference of dynamic resource allocation strategies, especially away from balanced growth, after a nutrient upshift or downshift [41]. The maturation constant for the reporter used in this study, along with the estimation of the degradation constant, was determined in previous work [41]. Similar to the estimation of growth rates, the inference of resource allocation strategies requires regularization parameters that were determined from the noise properties of the fluorescence intensity measurements for each experiment. As a side product, the inference method also returns an estimate of the total protein concentration, that is, the sum of the concentrations of the mature (observed) and immature (non-observed) protein.

**Figure 2:**
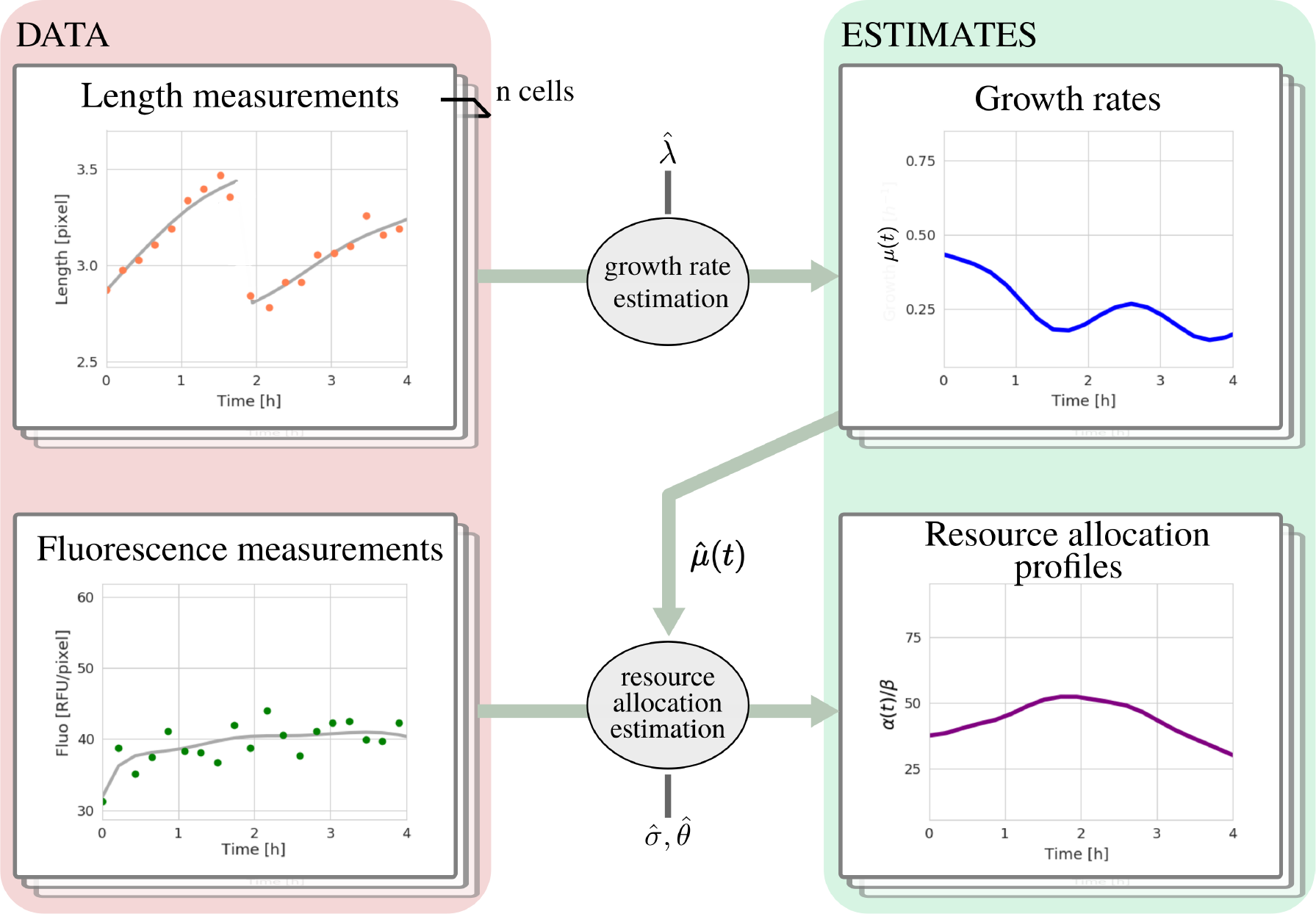
Inference procedures for estimating growth rates and resource allocation strategies from single-cell data. Time-lapse measurements of the length of mother cells are used as input by the growth-rate estimation method, returning time-varying estimates of the growth rate *µ*(*t*). The growth rate estimates 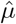, along with time-lapse fluorescence intensity measurements, corresponding to intracellular ribosomal concentrations, are then used for the estimation of single-cell resource allocation strategies *α*(*t*)*/β*. The two estimation methods rely on regularization to cope with measurement noise and return smoothed estimates. This requires values for the regularization parameters (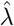 and 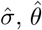) which are estimated for each experiment to account for possibly different experimental conditions, machine settings, etc. (*Materials and methods* and Text S2).

Fig. 1E shows the resource allocation strategy estimated for a single cell and Fig. 1D the corresponding fit to the fluorescence data for this cell. The figure also displays the mean resource allocation strategy estimated for all cells in the experiment. We observe that *α*(*t*)*/β* immediately increases after the upshift from acetate to glucose and reaches a new, higher steady-state level quite fast. The higher value of *α*(*t*)*/β* after the upshift means that a larger fraction of the protein synthesis rate is allocated to the ribosomes during fast growth on glucose, consistent with the growth law stating that a higher growth rate requires a higher ribosome concentration [8, 9]. For the downshift, the dynamics of adaptation to acetate are slower and more complex, involving a transient undershoot of *α*(*t*)*/β* lasting approximately five hours. This undershoot has not been observed before, but previous experiments did not follow (ribosomal) proteins with the same sampling density and over the same long time interval as this study [23, 24]. Interestingly, the undershoot is accompanied by a (less pronounced) undershoot in growth rate, which was observed in previous work as well [39].

### Variability of ribosomal resource allocation during balanced growth

During balanced growth, we expect the ribosome concentration to reach steady state, which implies *r*(*t*) = *α*(*t*)*/β* (Eq. 1). As explained above, the inference method of Fig. 2 allows us to estimate both the (total) ribosome concentration *r*(*t*) and the (scaled) resource allocation strategy *α*(*t*)*/β* from the data. In order to verify if these two terms agree, we computed during balanced growth on glucose and acetate (more than 5 generations after an upshift or downshift), for every individual mother cell, the average ribosome concentration and average resource allocation strategy over each generation. The results, consisting of more than 800 data points for individual generations of individual cells are shown in the scatter plot of Fig. S7. The two quantities are strongly correlated (*R*^2^ = 0.86), confirming that the method is capable of recovering the direct relation between ribosome concentration and resource allocation at steady state.

The steady-state data enable another straightforward test. The growth law for ribosomes, the linear increase of the ribosome concentration with the growth rate, has been extensively studied for populations of bacteria grown in different media. Nothing is known, however, about the correlation of the ribosome concentration with the growth rate in individual cells grown in the same medium. While one might expect a linear relation in this case as well, the test of another population-average growth law on the single-cell level shows that some caution is warranted. Taheri-Araghi *et al*. [32] found strong deviations from the expected exponential dependence of newborn cell volume on growth rate for individual cells grown in the same medium.

We first verified that the average of the growth rates and ribosome concentrations taken over individual generations of individual cells growing on acetate or glucose is consistent with the growth law established from population-level measurements (Fig. 3A). We then plotted the individual growth rates and ribosome concentrations in each of the two media (Fig. 3B). As can be seen, there is only a weak positive dependence of ribosome concentration on growth rate for growth on glucose and acetate. In both cases, there is a huge variability in growth rates and ribosome concentrations, such that individual cells can grow fast with low ribosome concentrations or slow with high ribosome concentrations. The same results were found in an independent replicate experiment (Fig. S8).

**Figure 3:**
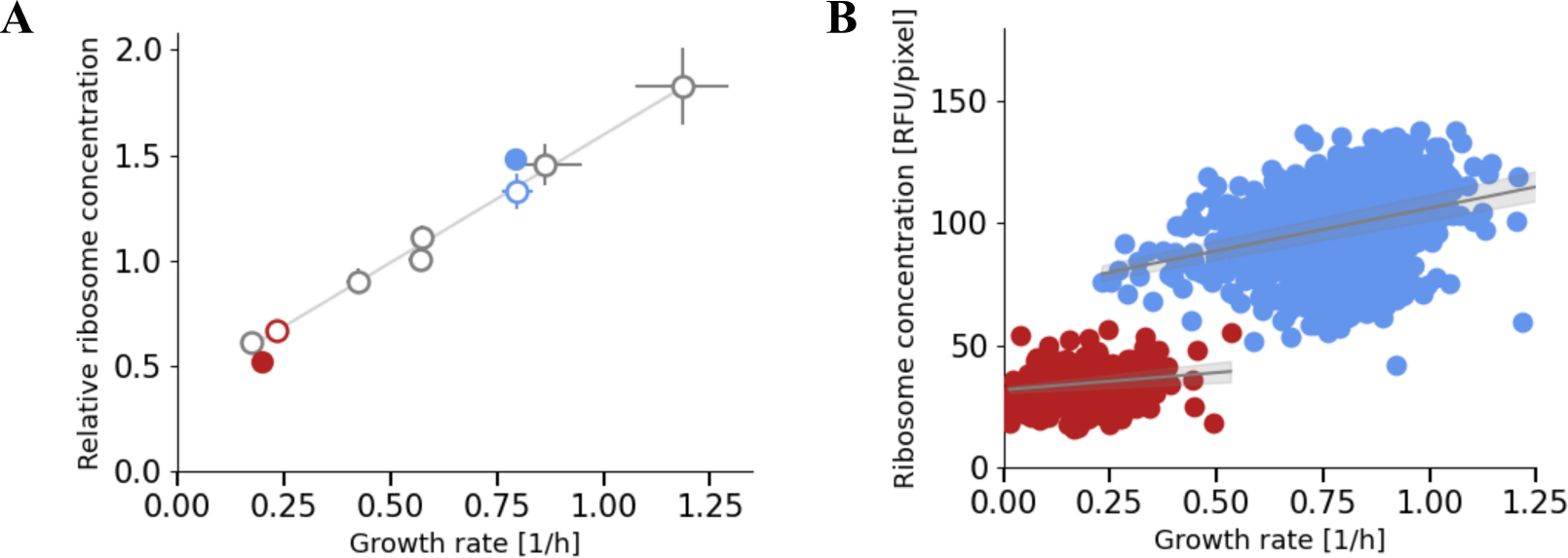
Single-cell resource allocation for ribosomes during balanced growth. **A**. Relation between the mean of growth rates and total ribosome concentrations for individual generations of individual cells of the Rib strain during balanced growth on acetate or glucose (filled circles). These results from the mother machine experiment in Fig. 1 are compared with population-level measurements for the same strain growing in batch on glucose, acetate, or other carbon sources (open circles) (Fig. S2). Data points for acetate are colored in red, for glucose in blue, and for other carbon sources in gray. In order to make the two data sets comparable, all ribosome concentrations were normalized by the mean of the average ribosome concentrations on glucose and acetate. A line was fitted to the batch data and plotted as a visual aid. Each point is the mean of 5 replicates and confidence intervals are given by two times the standard error of the mean. The growth rates and the relative ribosome concentrations during growth on glucose and acetate agree between the batch growth and microfluidics datasets. **B**. Relation between ribosome concentrations and growth rates for individual generations of individual cells during balanced growth on glucose (blue) and acetate (red). A line was fitted to each dataset and plotted in gray, along with the confidence bands representing two times the standard deviation. We observe weak dependence of the ribosome concentration on the growth rate on the single-cell level (34.9 *±* 2.9 for glucose and 14.2 *±* 5.7 for acetate), contrary to the population-average growth law for ribosomes shown in panel A.

How can this deviation from the population-level growth law be explained? The sum of the growth rate and the degradation rate is related to the time-varying total protein synthesis rate by *µ*(*t*) + *γ* = *β v*_*prot*_(*t*) (Text S1). That is, for the total protein concentration to remain constant, the rate of newly synthesized proteins *v*_*prot*_(*t*) must match, at every time-point, the sum of the rate of protein decay by growth dilution or degradation: (*µ*(*t*) + *γ*) (1*/β*). The protein synthesis rate can be defined as the product of the concentration of actively translating ribosomes *r*_*act*_ and the translation elongation rate *σ*, i.e., *v*_*prot*_(*t*) = *r*_*act*_(*t*) *σ*(*t*), where active and inactive ribomes make up the total ribosome pool: *r* = *r*_*act*_ + *r*_*inact*_ [25]. Careful measurements have shown that, in the conditions considered here, on average about 10% of ribosomes in the *E. coli* cell are inactive [45], and thus form an unused ribosome reserve [25, 34, 46].

Combining the above equations, we find

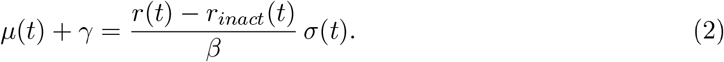

This relation provides a possible explanation for the observed lack of correlation between ri-bosome concentration and growth rate at steady state. It suggests that the pool of inactive ribosomes and/or the translation elongation rate vary over individual cells and across generations. This high variability could be due to, for example, active regulation of ribosome availability or to passive regulation of the elongation-rate by the size of amino acid and ATP pools [45-47].

The lack of correlation between growth rate and ribosome concentration is surprising in the light of the hypothesis, exploited in mathematical models [11, 14, 15, 18-20, 48, 49], that bacteria have evolved to optimize their growth rate. One would expect growth-rate maximizing cells to tightly control their investment in costly ribosomal machinery and avoid wasting resources on inactive protein synthesis capacity. While suboptimal from the perspective of growth-rate maximization during balanced growth, the maintenance of a ribosome reserve may be beneficial in other ways. It has been suggested, for example, that a ribosome reserve can be exploited for rapid adaptation to environmental changes [25, 26, 34, 46, 50-52].

### Fast and slow adapters after a nutrient upshift

In order to test the possible role of the ribosome reserve in speeding up the adaptation of resource allocation after a change in environment, we reconstructed the adaptation trajectories following a nutrient upshift or downshift. Note that during balanced growth it holds by Eq. 1 that *r* = *α/β*, so the (total) ribosome concentration can be used as a proxy for the fraction of protein synthesis capacity allocated to the synthesis of ribosomes in that condition. This is no longer true during a nutrient upshift or downshift, when the dynamics of ribosome concentration and resource allocation may temporarily diverge.

We first computed the average over all mother cells of the estimated adaptation dynamics of *µ* and *α/β* for an acetate-glucose upshift and plotted the trajectory (Fig. 4A). As can be seen, the trajectory has two consecutive phases. In the first phase after the upshift, resource allocation jumps to a high value, which then remains approximately constant while growth rate gradually increases. In the second phase, *µ* and *α/β* jointly increase, along the line of the growth law, towards the steady-state value for balanced growth on glucose. The corresponding time-courses for *µ* and *α/β* are shown in Fig. 4D,G. A sharp increase in resource allocation occurs immediately after the upshift, but it takes more than 6 hours to reach the steady-state value for balanced growth on glucose. For the growth rate, the increase towards the steady-state value is more gradual over the whole adaptation period.

**Figure 4:**
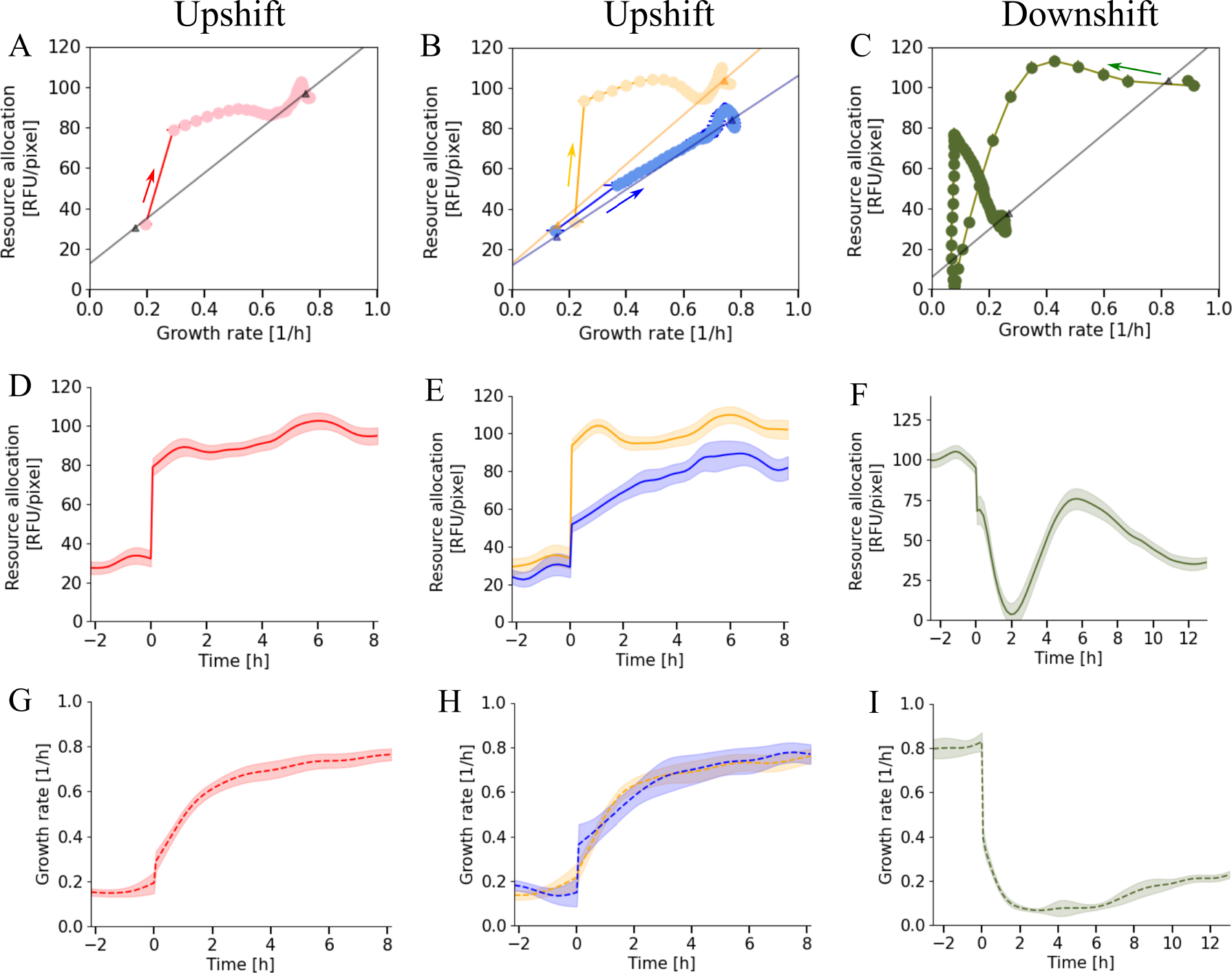
Adaptation dynamics of growth rate and ribosomal resource allocation after nutrient upshift and downshift. **A**. Mean adaptation trajectory of the ribosomal resource allocation strategy *α/β* and the growth rate *µ* after an acetate-glucose upshift applied to the Rib strain growing in a mother machine. The resource allocation strategies and growth rates were estimated from the data using the inference methods of Fig. 2 and averaged over 129 cells. The arrows indicate increasing time after the upshift. The triangles denote the average values of *α/β* and *µ* during balanced growth on acetate (before the upshift) and glucose (after the upshift). The latter values were determined by computing for each cell the mean growth rate over a period of balanced growth (*>* 2 h), and then averaging these values over the individual cells. The black line through the population average before and after the upshift is shown as a visual aid. The trajectories show that the adaptation of resource allocation and growth rate are uncoupled in a first phase and coupled in a second. **B**. Clustering of the single-cell resource allocation strategies for ribosomes after the upshift using k-means (*Materials and methods*) reveals two distinct trajectories for fast adapters (orange, 4 cells) and slow adapters (blue, 45 cells). **C**. As in panel A, but for a glucose-acetate downshift carried out in the same experiment (130 cells). The average adaptation trajectory displays an undershoot of resource allocation before settling at the steady-state value for growth on acetate. **D-F**. Time-courses of the resource allocation strategies in panels A-C, respectively. **G-I**. Time-courses of the growth rates in panels A-C, respectively. Confidence intervals for resource allocation strategies and growth rates are given by two times the standard error of the mean.

What is the cell-to-cell variability of the resource allocation strategies and growth rates underlying the averaged adaptation trajectory in Fig. 4A? We performed a clustering analysis to classify the resource allocation strategies of the individual cells in the first hours after the upshift (Fig. S9). We found that grouping the data into two clusters provided a statistically sound and biologically informative view of the variability in the data (*Materials and methods*). The adaptation trajectories for these two clusters are plotted in Fig. 4B. Interestingly, the two-phase trajectory of the population average turns out to be the superposition of the trajectories for the individual clusters. A first cluster of fast adapters (84 of 129 cells) immediately jumps to the resource allocation value for balanced growth on glucose and approximately remains at this value when increasing the growth rate. A second cluster of slow adapters (45 of 129 cells) mostly increases growth rate and resource allocation simultaneously, along the line of the growth law. The time-courses of the growth rates and resource allocation strategies for the two clusters underline these observations (Fig. 4E,H). This analysis of resource allocation strategies on the single-cell level is highly reproducible across replicate experiments (Figs S10 and S11).

The cluster of fast adapters confirms the hypothesis, proposed by Bremer and Dennis [24], that after an upshift cells immediately set resource allocation to the steady-state value on glucose and then gradually adapt their growth rate. A recent kinetic model by Erickson *et al*. [23], which includes feedback regulation of *α* by the translational activity *σ*, made a very similar prediction. These models describe only part of the observed dynamics, however, as the cluster of slow adapters gradually increase both the resources allocated to ribosomes and the growth rate, by following the linear relation of the growth law. The data do not agree with the hypothesis that the adaptation trajectory maximizes biomass accumulation, which was predicted to give rise to an oscillatory pattern for *α* [18].

What causes the population to divide in fast and slow adapters? A possible clue can be obtained by considering the allocation of resources to ribosomes right before the upshift (Fig. 4E).

In this experiment and in the replicate experiments (Figs S10 and S11), the mean value of *α/β* is higher in the cluster of fast adapters than in the cluster of slow adapters. By the steady-state relation *r* = *α/β*, this must also hold for the mean value of the ribosome concentration, as confirmed by Fig. S12. A higher ribosome concentration before the upshift enables the existence of a higher ribosome reserve [25, 26], and may thus favour a rapid reallocation of resources to ribosomes when the medium is switched from a poor to a rich carbon source. The consequent higher value for *α/β* after the upshift is predicted by the model of Eq. 1 to lead to a higher ribosome concentration (Text S3). This is confirmed by the experimental data: fast adapters maintain a higher ribosome concentration after the upshift (Fig. S21). The reasoning underlying this explanation of the occurrence of fast and slow adapters has been summarized in Fig. 5.

**Figure 5:**
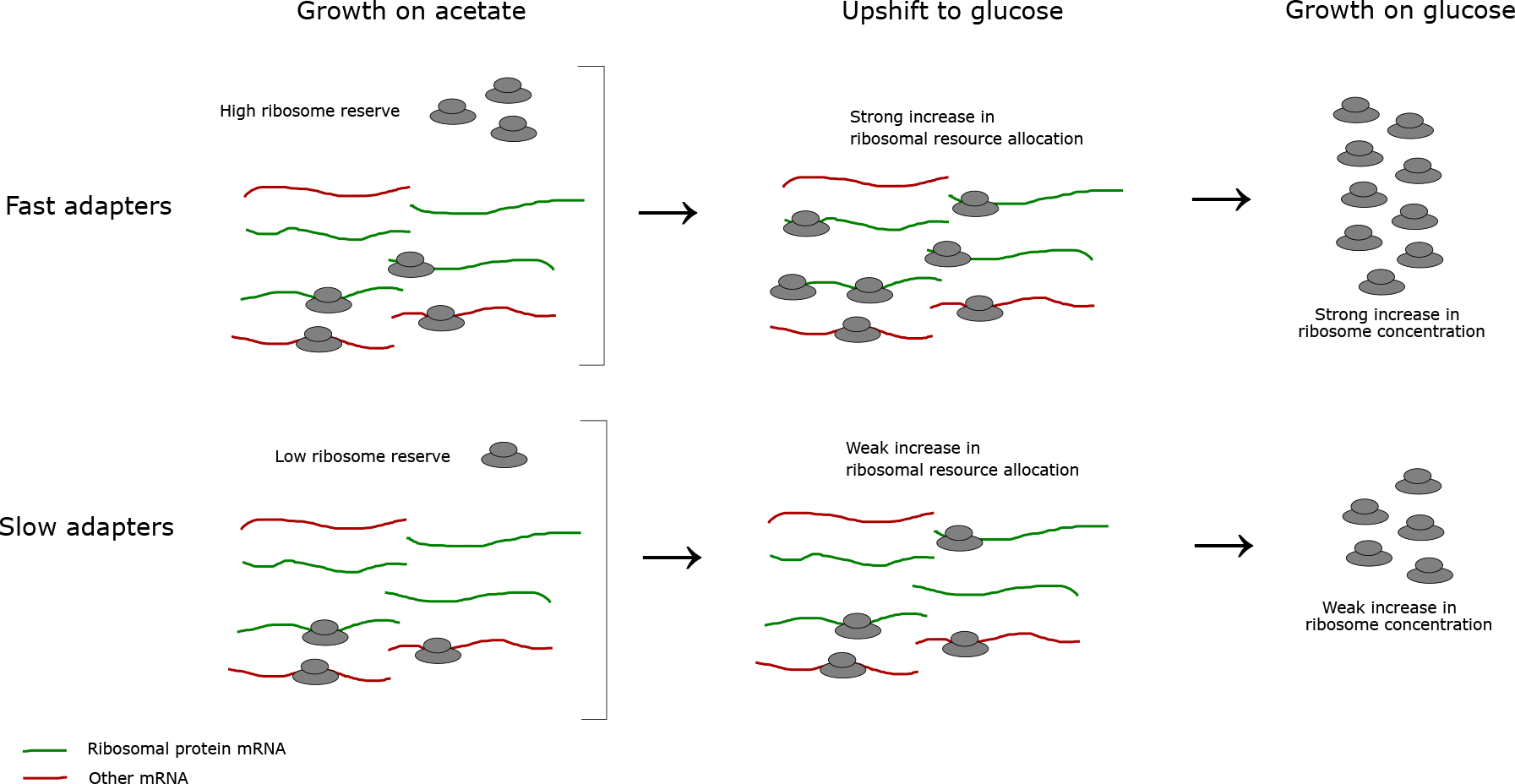
Explanation for the occurrence of fast and slow adaptation of resource allocation after a nutrient upshift. Fast adapters have a higher ribosome concentration before the upshift from acetate to glucose (Fig. 4E and Fig. S12). This presumably allows them to have a higher ribosome reserve, that is, a higher excess capacity that can be allocated to the synthesis of new ribosomes immediately after the upshift. This allows fast adapters to make a higher jump in ribosomal resource allocation after the upshift (Fig. 4E), as illustrated in the figure above by the increased proportion of ribosomes translating ribosomal mRNA. The higher allocation of ribosomes to the synthesis of new ribosomes in fast adapters leads to a stronger increase in the concentration of ribosomes (Fig. S21). Faster adaptation of resource allocation does not necessarily lead to faster adaptation of growth rate (Fig. 4H and Fig. S22).

One expects that, on average, cells that rapidly adapt ribosomal resource allocation after the upshift have an increased capacity to step up their growth rate. Previous work proposed a dynamic growth law [25] relating the difference in ribosome concentrations during balanced growth on a rich and a poor carbon source to the magnitude of the jump in growth rate directly after the upshift. Like for the classical growth law during balanced growth, the population-averaged data from the microfluidics experiments are consistent with the predictions of the dynamic growth law (Fig. S22). For the individual cells, however, the magnitude of the jump in growth rate directly after the upshift is not correlated with the difference in ribosome concentrations during growth on glucose and acetate. While the estimation of jumps in growth rates directly after the upshift is inherently noisy for individual cells, inspection of growth-rate adaptation averaged over the clusters of fast and slow adapters points in the same direction (Fig. 4H). Fast adapters do not grow at a higher rate than slow adapters during the upshift, as might have been expected from their higher ribosome concentrations. The replicate experiments do not provide conclusive evidence for a growth advantage of fast adapters either (Figs S10 and S11).

We performed the same analysis of ribosomal resource allocation strategies for a glucose-acetate downshift. We observed a complex pattern of adaptation, during which both *µ* and *α/β* sharply drop directly after the downshift, before partially recovering and eventually settling at the lower growth rate and ribosomal resource allocation characteristic for growth on acetate (Fig. 4C,F,I). In this experiment, resource allocation dropped just before growth rate, but in a replicate experiment carried out under identical conditions, the time-ordering of events was the inverse (Fig. S11). The differences in time-ordering may be due to heterogeneity in the drop of resource allocation among the individual cells in the population (Fig. S13). It is difficult to reach a decisive conclusion, however, given that the sampling density was too low to reliably capture events occurring in such rapid succession.

## Discussion

Growth laws empirically describe the resource allocation strategies adopted by microbial cells [4, 53]. These relationships have been established for many species and experimental conditions, yet their scope has been mostly limited to balanced growth. A few studies have focused on growth in unbalanced conditions, especially the adaptation of resource allocation after a sudden change in nutrient availability [18, 23, 24], but experimental data remain scarce. Moreover, the available data quantify adaptation on the population level, thus ignoring possible heterogeneous behavior of the individual cells making up the population [27-33]. The question that remains unanswered is therefore how resource allocation strategies vary across the cells in an isogenic population, both during balanced and unbalanced growth.

In this study, we addressed the above question by quantifying, for the first time, ribosomal resource allocation strategies on the single-cell level. We created fusions of a ribosomal subunit with different fluorescent reporters in *E. coli* and conducted time-lapse microfluidics experiments in which we recorded the dynamic adaptation of the abundance of the tagged ribosomes after a sudden change in carbon source (Fig. 1). Moreover, we developed robust statistical inference methods to reconstruct resource allocation strategies from the resulting fluorescence data, using a mathematical model taking into account protein synthesis, maturation, degradation, and growth dilution (Fig. 2). The methods are generic and applicable to the inference of resource allocation strategies in other microorganisms and experimental scenarios, not only for ribosomes but for any protein tagged with a fluorescent reporter.

We first explored the relationship of the growth rate and the allocation of resources to ribo-somes during balanced growth on glucose or acetate (Fig. 3). Whereas the averaged data agree with the well-established growth law for ribosomes, which posits a linear dependence of the ribo-some concentration on the growth rate across different media, we discovered that the correlation is almost negligible in the case of individual cells growing in the same medium. The absence of a correlation between growth rate and ribosome concentration indicates that ribosome activity is highly variable across a population of *E. coli* cells. This agrees with the observation made for another population-level growth law, relating growth rate to cell volume immediately after cell division [32]. A recent theoretical study also predicted that, in the conditions considered here, fluctuations in enzyme rather than ribosome concentrations account for the variability of growth rates in individual cells [54].

The observed variability of ribosome concentrations during balanced growth may be an un-avoidable consequence of the stochasticity of gene expression [55-57], without a specific biological function. In particular, the synthesis of ribosomes in *E. coli* is regulated by the signaling molecule guanosine tetraphosphate (ppGpp) [47, 58-61]. The number of molecules of ppGpp may diverge across the cells in a population, due to heterogenous activity of the enzymes responsible for ppGpp synthesis and hydrolysis. As a consequence, *E. coli* may experience fundamental limits on the capacity to precisely regulate resource allocation on the single-cell as compared to the population level. The large difference in ribosome contents of the individual bacteria, up to three-fold, is nevertheless surprising. Ribosomes are costly, abundant protein complexes, highly conserved in evolution, and responsible for the synthesis of proteins, the major component of biomass. Moreover, a high ribosome concentration in a mother cell tends to be preserved over more than 10 generations (Fig. S21), which would not be expected if the observed variability was due to random gene expression noise alone.

Alternatively, the variability of gene expression may have a biological function, as demon-strated in various examples of bacterial stress responses [27, 28, 31, 62]. The maintenance of a ribosome reserve in an isogenic population could have a fitness advantage, by trading off maximal growth during balanced growth against the capacity to quickly react to an environmental change [25, 26, 34, 46, 50-52]. If *E. coli* had adopted such a bet-hedging strategy, one would expect bacteria with a higher ribosome concentration during growth on acetate, and therefore a likely higher ribosome reserve, to outgrow cells with a lower ribosome concentration when the medium is switched to glucose. This is not observed in our data. Whereas a higher ribosome concentration before the upshift leads to faster adaptation of resource allocation during the upshift (Fig. S12), this does not systematically translate into higher growth rates (Fig. 4 and Figs S10 and S11). It is possible though that fast adaptation of resource allocation enabled by the ribosome reserve confers a growth advantage in other conditions, for example a change in environment requiring *de-novo* expression of specific necessary enzymes [31, 51].

In conclusion, the present work identifies, for the first time, an important source of cellular heterogeneity on the level of ribosomes, the molecular machines at the heart of cellular self-replication. This raises interesting questions on whether *E. coli* cells have evolved to exploit or tolerate the variability in ribosomal resource allocation [63, 64].

## Materials and methods

### Strain construction

The strain used in this study were derived from the *E. coli* K12 strain BW25113 [36] where the *fhuA* gene was deleted by homologous recombination to confer resistance to phage infections [65]. To visualise the ribosomes, we used a construction similar to the one described by Bakshi *et al*. [35] where the gene coding for the green fluorescent protein GFPmut2 [49] was fused to the C-terminus of *rpsB*, the gene coding for the S2 ribosomal subunit, via a flexible linker of three amino acids (LEI).

The cloning procedure consisted in first amplifying a PCR fragment carrying 50 bp of sequence identity with the target insertion site. The PCR fragment, called (positive-negative selection) “cassette”, contains two divergently transcribed genes: the gene coding for resistance to kanamycin, transcribed from a constitutive promoter, and the gene coding for the toxin CcdB, transcribed from the arabinose-inducible promoter pBAD [66]. This fragment was recombined into the target site using the pSIM plasmid that provides the lambda Red recombination functions [67]. The successful insertion was selected on LB containing glucose and kanamycin. Glucose prevents expression of the CcdB toxin and kanamycin selects for the presence of the cassette on the chromosome. In a second recombination step, the cassette was removed by recombination of a PCR fragment containing the sequence to be integrated flanked by the same 50 bp homology regions to the target site. Successful recombination is selected on LB containing 1% arabinose. Arabinose activates the pBAD promoter, and therefore the expression of the toxin CcdB, if the cassette is still present on the chromosome. A successful recombination will have removed the cassette and thus allow growth of the strain in the presence of arabinose. The resulting strain was called Rib, for ribosomal reporter.

The strain used in this study is defined in Tab. S1 and graphically presented in Fig. S14. The construction was verified by sequencing of the modified chromosomal region. Primers are available upon request.

### Growth media

For all experiments, we used M9 minimal medium [68] supplemented with vitamin B1 (5 mg/L) and mineral trace elements. Pre-cultures for the microfluidics experiments were prepared in the above medium supplemented with 2 g/L of acetate and 0.1 g/L of bovine serum albumin (BSA). The medium used for the microfluidics experiments was supplemented with 2 g/L of acetate or glucose. The experimental conditions of the individual microfluidic experiments are detailed in Tab. S2.

For strain validation in batch experiments, minimal medium was supplemented with 2 g/L of one of the following carbon sources: D-glucose, acetate, D-xylose, pyruvate, glycerol, D-fructose, D-galactose and maltose. Furthermore, 2 g/L of casamino acids were added into D-glucose and glycerol in separate conditions. For pad experiments, the medium was supplemented with 2 g/L of acetate before adding agarose.

### Batch growth experiments

All strains were streaked from the freezer stock on LB agar plates several days prior to the experiments. Overnight cultures were grown by inoculating a single colony in liquid minimal medium supplemented with specified carbon sources. Bacterial growth for all experiments occurred at 37°C and liquid cultures were shaken at 200 rpm.

Overnight cultures of the wild-type and reporter strain were diluted into a 96-well microplate at an OD600 of 0.02 where each well contained the same medium as used for the preculture. 2 mm glass beads were added in each well to improve oxygenation. The bacteria were left to grow for up to 24 hours at 37°C in a Tecan Infinite 200M plate reader, while monitoring the absorbance and green fluorescence.

Outliers in the absorbance and fluorescence curves due to the light reflection by the beads were filtered using WellInverter [69]. The fluorescence background was corrected by subtracting the fluorescence emitted by the wild-type strain in the same conditions. The absorbance curves were corrected by subtracting the background of a well containing medium but no bacteria. The growth rate during balanced growth of the population was estimated directly from the absorbance curves, using the method described by Zulkower *et al*. [70]. In order to obtain a proportional estimate of the reporter protein concentration during balanced growth, the background-corrected fluorescence curves were divided by the background-corrected absorbance curves.

### Microfluidics experiments

#### Fabrication of microfluidic devices

The master-molds were obtained from Epsilon Micro Devices (College of Engineering, Zahn Center, 5500 Campanile Drive, San Diego CA). In order to produce PDMS mother machines, we adapted the protocol from the mother machine handbook published by the Jun laboratory at UC San Diego (https://jun.ucsd.edu/files/mothermachine/mothermachinehandbook.pdf). In particular, we mixed a curing agent and polymer base at a 1:10 ratio (Sylgard Elastomer Kit 184). The mix was poured into the mold and trapped air bubbles were removed in a vacuum chamber. The resulting devices were cured for a minimum of 2 hours at 65°C. Devices were then removed from the master-mold and a 0.75 mm punch was used to create the inlet and the outlet. Before use, the devices were treated with pentane and acetone and were left to dry overnight. A plasma cleaner (PDC-32G Harrick Plasma) operated at high intensity for 40 seconds was used for surface activation of the devices and previously cleaned microscope slides. Each device was placed on a microscope slide and then put at 65°C for 10 minutes. 50 mg/mL of BSA was injected into the sealed devices for surface passivation and the devices were then incubated at 37°C for 1 hour.

#### Device loading and medium shifts

Precultures were performed in the same way as for the batch growth experiments described above. The optical density of strains growing in minimal medium with acetate was monitored and the cells were harvested during balanced growth (after at least 6 generations). The cells were then washed with fresh medium containing 50 mg/mL of BSA, concentrated around 100-fold and loaded into the device. The individual cells growing in the microchannels were left overnight in acetate to reach balanced growth before starting image acquisition.

An Elveflow pressure controller (OB1 MK3) equipped with a microfluidic flow sensor (0 to 50 *µ*L/min) was used to assure a constant flow delivery of 20 *µ*L/min to the device. The bottles of medium to be changed at the time of medium switches were prepared and put at 37°C many hours before the switch. After the switch, 2 to 3 minutes were required for the new medium to reach the device. The corresponding dead-volume and dead-time were taken into account in all our analyses. Fig. S3 illustrates the scenario for a typical microfluidics experiment.

#### Image acquisition

A Zeiss Axio Observer Z1 inverted microscope equipped with a halogen lamp controlled by a Vincent D1 shutter and a motorized xy-stage was used to perform the experiments. The focus was maintained by Zeiss Definite Focus. Images were recorded using a Zeiss Plan-APOCHROMAT 63x/1.40 oil objective and a Digital CMOS camera (Hamamatsu Orca-Flash 4.0). A constant temperature of 37°C was ensured by a Peltier-equipped box with temperature sensors. The setup was controlled by the Micromanager software [71] and images were recorded using the Multi-Dimensional Acquisition (MDA) feature. A phase contrast image (70 ms exposure) was recorded along with a (green) fluorescence image every 11-13 minutes for multiple positions in the device. To obtain fluorescence images, a Zeiss Colibri 7 was used equipped with LEDS fixed at 6% intensity. The following filter, purchased from Chroma, was used: GFP (49002).

### Pad experiments

Agarose pads were prepared by melting medium supplemented with appropriate carbon sources and 1.5% of low-melt agarose. 1 mL of hot mix was placed between two 2x2 cm microscope cover slips and let to solidify for one hour in a Petri dish. The upper cover slip was then removed and the pad was cut into four distinct pieces to accommodate multiple strains. Overnight cultures of the reporter strain and the wild-type strain were diluted to an OD600 of 0.1, and 2 *µ*L of each culture was spotted onto a distinct section of the pad. The pad was left to dry for 15 minutes and a microscope slide was then placed on top of the pad. The pad was sealed using paraffin and transferred to the microscope. The same image acquisition setup and parameters were used as described for the mother machine experiments.

In total, two pad experiments were conducted in order to quantify the autofluorescence of our reporter strain as compared to the autofluorescence of the wild-type strain. The obtained images were analyzed in ImageJ and 70 cells were manually segmented using the phase-contrast channel. The fluorescence intensity was then extracted for each segmented cell, fluorescence channel and frame. For each cell, the fluorescence intensities were averaged across all frames to obtain a more robust estimate of the autofluorescence.

### Image processing

#### Pre-processing and segmentation

For image pre-processing and segmentation, we used the mother machine segmentation software tool BACMMAN [38]. Raw images were sorted and imported into the software. After a pre-processing step in which the images were stabilized and de-noised, the microchannels and individual bacteria were segmented on the green fluorescence channel. For each experiment, empty microchannels and microchannels presenting double-loading were discarded from the analysis, along with a few cells that stopped growing at various phases of the experiment. The segmentation was performed using the BacteriaFluo algorithm. In our study, we included around 350 mother cells (Tab. S2). After segmentation, the mother cells and first-generation daughters were visually checked and the few segmentation errors (*<* 5%) manually corrected.

#### Post-processing

For each segmented cell, we obtained the mean fluorescence intensity RFU/pixel, the length of each cell pixel and various other statistics containing information on position and lineage. For the chosen settings, one pixel corresponds to 0.11 *µ*m. These results were imported into Python where several post-processing steps were carried out. The camera noise and background were corrected for each channel by subtracting the base value we obtained with a closed shutter.

The autofluorescence was evaluated using pad experiments, as described above. The autofluorescence of the wild-type strain was found to be around zero, after background correction, and negligible in comparison with the fluorescence emitted by the reporter strains (Fig. S15). We therefore applied no autofluorescence correction to the measured fluorescence intensities.

To detect any reporter photobleaching effects in our experiments, we performed a microfluidics control experiment with the reporter strain growing in balanced growth in minimal medium with acetate (Tab. S2). The same image acquisition parameters as described above for the microfluidics experiments were used. The images thus obtained were segmented and the fluorescence intensity extracted (Fig. S16). The curve was fitted by a straight line and the obtained slope coefficient was found to be indistinguishable from zero, so no photobleaching correction was applied in the measurement models.

### Model definition and calibration

To reconstruct the resource allocation profiles from single-cell fluorescence data, we developed mechanistic models that take into account fluorescent protein maturation, degradation and growth dilution.

GFP has first-order maturation kinetics [72], so the dynamics of the RpsB-GFPmut2 fusion in the Rib strain can be described by the following simple model (Text S1 and [41]):

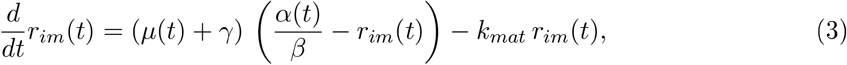

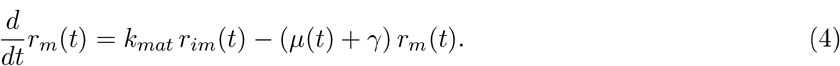

*r*_*im*_ and *r*_*m*_ refer to the concentrations of ribosomes tagged with immature and mature GFP, respectively, in units RFU/pixel. Maturation is characterized by the constant *k*_*mat*_ 1/h. Protein degradation is determined by the degradation constant *γ* 1/h. *α*(*t*)*/β* is the (scaled) dynamic resource allocation strategy for ribosomes in the cell that can be reconstructed from single-cell fluorescence data using the methods described below. Note that the sum of Eqs 1-2 results in Eq. 1 in the main text, describing the dynamics of the total (mature and immature) ribosome concentration.

The maturation and degradation parameters for the GFPmut2 reporter were determined experimentally in previous work [41], in experiments where an antibiotic (Chloramphenicol) was added to a growing culture to stop translation and the ensuing fluorescence was measured. The parameter values are summarized in Tab. S3.

### Growth-rate estimation

Growth rates of mother cells over the entire duration of an experiment were inferred from measured cell lengths by means of a custom-made estimation method. The method is based on the following model of growth rate *µ*(*t*):

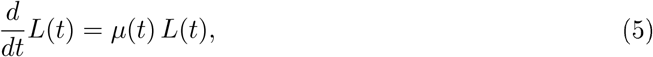

where *L*(*t*) is the time-varying length of a mother cell, expressed in pixel units. Cell length is a good approximation of cell size in rod-shaped bacteria like *E. coli* [2, 37]. Contrary to many growth-rate estimation methods used for microfluidics data, we did not assume that growth rates are constant between two consecutive cell divisions. However, growth rates were assumed constant between two consecutive measurement times, which is reasonable given the sampling density (once every 11-13 min, corresponding to 4-30 length measurements per generation).

As explained in Text S2, the above assumptions result in an estimation problem defined for a piecewise-linear model with as many equations as cell length measurements. Because the solutions of this problem are underdetermined by the (noisy) experimental data, we employed a regularized least squares method [73]. Regularization involves a cost function penalizing rapid changes in growth rate, multiplied with a regularization parameter *λ >* 0. We fixed an appropriate value for *λ* by means of cross-validation [74]. In particular, we estimated an optimal value 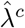 for every mother cell *c* considered in an experiment. The median of these estimates was used as our best choice for *λ*, which makes the latter robust to measurement outliers caused by a variety of experimental errors, such as an occasional loss of focus. In order to allow for rapid changes in growth rate after nutrient upshifts and downshifts, no regularization was enforced immediately after a growth transition (Text S2). All of the above-mentioned computations were carried out in Python 3.6, using the code available in File S1.

The results of the growth-rate estimation method were validated in two different ways. First, we assessed the performance of the method by applying it to synthetic cell length data generated by means of Eq. 5 for a given time-course growth-rate profile. To the cell lengths thus generated, we added Gaussian noise in agreement with the experimentally observed distribution. From these synthetic data, the method succeeded in robustly reconstructing the known growth-rate profile (Fig. S17). Second, we compared the performance of the method with a baseline approach consisting in the fit of a linear curve to log-transformed length data between two consecutive cell divisions, corresponding to the assumption of a constant growth rate during each generation time. During balanced growth, when the growth rate is approximately constant, the two methods give comparable results. However, when the assumption is not satisfied, especially during transitions between growth phases, our method gives more plausible estimates (Fig. S6).

### Inference of resource allocation strategies

#### Kalman smoothing method

For each mother cell, the unknown resource allocation strategy *α*(*t*) was inferred from noisy fluorescence intensity measurements *y*_*k*_ = *r*_*m*_(*t*) + *∈*_*k*_, obtained at time-points *t*_*k*_, *k* = 1, …, *n*. The measurement errors *∈*_*k*_ form a sequence of independent random variables with mean zero and variance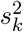. We assumed that all measurements obtained in a specific growth phase in an experiment, that is, in a specific growth medium, have the same variance. The variance was estimated from the fluorescence data when the cells had been in balanced growth in the medium for more than 6 generations, and *α*(*t*) can be assumed stationary. In particular, we determined the variance from the observed standard deviation around a constant mean in the corresponding time interval. In order to increase the robustness of the estimate, we selected the median of the values for the individual mother cells.

Estimation of the unknown resource allocation profile demands the formulation of a probabilistic prior on *α*(*t*). This prior was expressed in the form of a stochastic differential equation

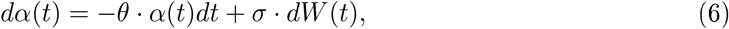

where *W* (*t*) is the standard Wiener (white noise) process, and *σ, θ* > 0 are parameters defining the (magnitude and time-scale of) of fluctuations in the resource allocation strategy [41]. We assume that *α*(0) has a zero-mean Gaussian distribution with variance *σ*^2^*/*(2*θ*), which implies that the process is stationary. The parameters *σ* and *θ* play the role of regularization parameters, ensuring that the inference of *α*(*t*) is robust to measurement noise. We denote the optimal values for these parameters by 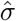 and 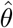 (see below).

With Eq. 6, the problem of calculating an optimal estimate 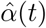 for *α*(*t*), given fluorescence intensity measurements *y*_1_, …, *y*_*n*_, becomes the Bayesian problem of calculating the conditional expectation

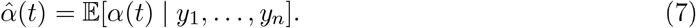

This reconstruction is optimal in the sense that it minimizes the variance of the estimation error at any time *t* 75. In order to solve the above problem, the measurement model and the stochastic prior were combined with the maturation models of the green reporter protein (Eqs 3-4), giving rise to the following linear stochastic differential equation model:

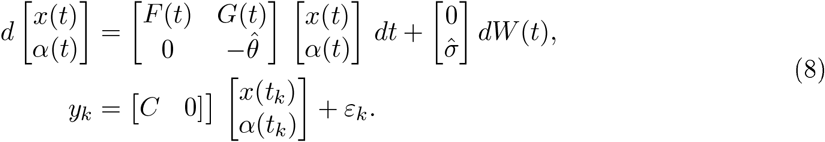

The vector *x* represents the state of the system, equal to [*r*_*im*_, *r*_*m*_]. Matrices *F* (*t*) and *G*(*t*) depend on the kinetic parameters in the maturation model, whose values are known (Tab. S3), as well as the estimated growth-rate profile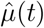, which was derived as explained above. More precisely, we have

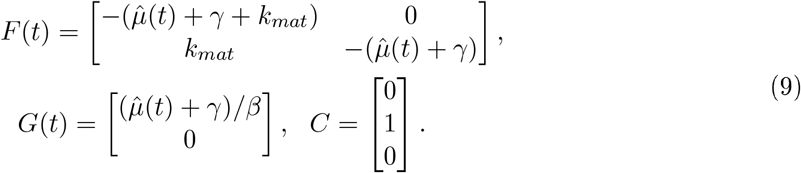

Recasting the system in the above form allowed the use of standard Kalman filtering and smoothing methods [76] to compute 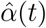 from the fluorescence intensity measurements. Since the value of *β* is unknown for our strain, we arbitrarily set it to 1. This means that in practice we estimate the scaled resource allocation strategy 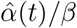 from the data, which allows us to monitor relative changes in resource allocation over time. Text S2 explains how Kalman filtering and smoothing were instantiated for this specific problem. Instead of solving the estimation problem for the entire measurement time-series, we treated the growth phases in different media separately, like for growth-rate estimation. At the medium switching times separating the growth phases, the resource allocation profile is expected to rapidly change over a short time interval, so that a smooth reconstruction of *α*(*t*) would be artifactual.

The computational complexity of the whole procedure is linear in the number of data points *n*, and is mostly determined by the inversion of the (small) matrices in Eq. 8. For a single mother cell with fluorescence time-series of *n* = 300 points (our case), the Python implementation using function `odeint` of the `scipy.integrate` module for numerical integration, requires 10 seconds. If the Gaussian assumptions are violated, the estimates computed by this procedure have the interpretation of optimal estimates (minimal error variance) in the class of linear functions of the data [75]. Thanks to this, the method is robust to moderate deviations from Gaussianity.

#### Automatic tuning of regularization parameters

An optimal choice of *σ* and *θ* was determined by solving the maximum likelihood problem

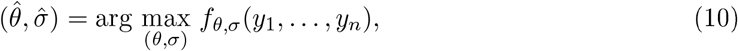

where *f*_*θ,σ*_(*·*) is the probability density function for the observed data under the stochastic prior of Eq. 6 [4]1. In practice, for any value of (*θ, σ*), evaluation of the above likelihood can be performed by Kalman filtering, in a similar way as for the estimation problem above. The details are given in Text S2.

The computational efficiency of Kalman filtering is important for the automatic tuning of the regularization parameters, because the optimization problem requires a large number of evaluations of the right-hand side of Eq. 10. Optimization was carried out numerically by the Python function `minimize` of the `scipy.optimize` (File S1). In a typical experiment with 100 mother cells and *n* = 300, the total computation time of the tuning procedure was around 2 hours.

For every mother cell *c*, we computed optimal regularization parameters 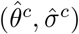 in the way described above. The final choice of the regularization parameters consisted in choosing the median of the values over all *c*. The latter value was then used for the estimation of the resource allocation strategy from the time-series data for every individual mother cell.

#### Validation of the method

The method was validated in two different ways. First, we generated synthetic fluorescence data using the model of Eq. 8, for a given resource allocation profile and noise characteristics corresponding to those of the real data. The Kalman smoothing estimation method was able to accurately reconstruct *α*(*t*) (Fig. S18). Second, we visually inspected the fit of the fluorescence intensities predicted by the model from the inferred resource allocation strategy and the actually observed fluorescence intensities for a large number of cells (Fig. 1). This was done both for the chosen optimal regularization parameters and (as a sensitivity test) for discarded non-optimal choices.

### Clustering of resource allocation strategies

In order to discover shared adaptation patterns across individual cells considered in an experiment, we clustered the reconstructed ribosomal resource allocation strategies after an upshift from acetate to glucose by means of a k-means clustering algorithm [77] implemented in Python (`sklearn` package [78]). Since all cells reach approximately the same steady-state levels of *α*(*t*)*/β*, we only clustered the data during the first two to three hours after the upshift (Fig. S9). In order to select the number of clusters in the k-means algorithm, we relied on the elbow method [79, 80]. The optimal number of clusters suggested by this method is 2 or 3 (Fig. S19). We decided to retain two clusters for the analysis, because as can be seen in the figure, they correspond to two clearly distinguishable patterns in the fluorescence intensities. Moreover, as shown in Fig. S20, the addition of a third cluster does not lead to distinctly different growth rate and resource allocation patterns. We checked that the mother cells in the clusters are homogeneously distributed over the positions in the microfluidics device.

### Predictions of dynamic growth laws

We experimentally tested the predictions of the jump in growth rate directly after the acetate-glucose upshift in the reference experiment, using the theoretical models of Mori *et al*. 25 and Korem Kohanin *et al*. [26].

Let 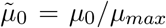 be the growth rate on acetate before the upshift (*µ*_0_), normalized with respect to the maximum growth rate of the *E. coli* BW25113 strain (*µ*_*max*_). Similarly, 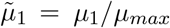 and 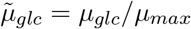 are the normalized growth rate directly after the upshift and the normalized growth rate on glucose that is eventually reached after the upshift, respectively. We used the following model (Eq. 7 in [26]):

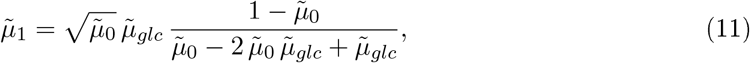

to predict 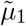 from values for *µ*_0_ and *µ*_*glc*_ from our data and a value for *µ*_*max*_ from the literature. In particular, we computed *µ*_0_ as the average value of the growth rate on acetate during 4 h preceding the upshift. Idem for *µ*_*glc*_, averaged over 2 h more than 10 generations after the upshift. For *µ*_*max*_, we used the value 1.6 h^*−*1^ for growth of the BW25113 strain in LB medium [81]. The experimental values for 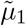 were obtained by dividing the growth rate directly after the upshift by the value of *µ*_*max*_. The model was applied both for values obtained for each individual mother cell and for values averaged over all mother cells in the experiment.

Let *ϕ*_0_ and *ϕ*_*glc*_ be the ribosomal protein fractions before the upshift and during balanced growth on glucose after the upshift, respectively. The following model (Eq. 8 in [25]) predicts the growth rate directly after the upshift:

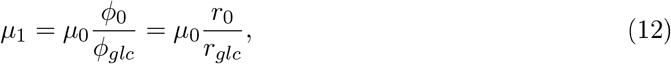

where we have replaced the ratio of ribosomal protein fractions *ϕ*_0_*/ϕ*_*glc*_ by the ratio of ribosome concentrations *r*_0_*/r*_*glc*_ before the upshift and during balanced growth on glucose after the upshift. If the total protein concentration remains approximately constant between these two time-points, then the two ratios are equivalent. We computed *r*_0_ and *r*_*glc*_ from the fluorescence data, by averaging the total ribosome concentration returned by the inference algorithm in the same time-intervals as *µ*_0_ and *µ*_*glc*_ above. Combined with the growth rate *µ*_0_, computed as above, this yields a predicted value for *µ*_1_. This prediction was compared with the experimentally determined value for *µ*_1_. Like for the previous model, the comparison was carried out for the values obtained for each individual mother cell and values averaged over all mother cells in the experiment.

## Data availability

All data supporting the findings of this study are available within the paper and its Supplementary Information.

## Code availability

The Python code to generate the figures in the main text and the Supplementary Information from the source data are available as File S1.

## Acknowledgments

This work was supported by the ANR project Maximic (ANR-17-CE40-0024). The authors would like to thank Grégory Batt and Jean-Luc Gouzé for comments on a previous version of the manuscript.

## Author contributions

AP, JG, and HdJ designed the study. AP, NG, and CP constructed the strains. AP and NG determined the experimental conditions. AP performed the experiments. IM and MVMG developed, implemented and supervised the fluorescence microscopy experiments in pads and mother machine. AP and EC developed and implemented the growth-rate and resource allocation inference procedures. AP, EC, JG, and HdJ analyzed the data. AP, EC, JG, and HdJ wrote the paper.

