## Supplementary material for "Single-cell data reveal heterogeneity of resource allocation across a bacterial population": SI figures

### Supplementary Tables, Figures, Movies and Files for "Single-cell data reveal heterogeneous resource allocation across a bacterial population"

Antrea Pavlou[1,2], Eugenio Cinquemani[1,2], Corinne Pinel[1,2], Nils Giordano[3], Mathilde Van Melle-Gateau[2], Irina Mihalcescu[2], Johannes Geiselmann[1,2,*,+], and Hidde de Jong[1,2,*,+]

[1] *Univ. Grenoble Alpes, Inria, Grenoble, France*
[2] *Univ. Grenoble Alpes, CNRS, LIPhy, Grenoble, France*
[3] *Nantes Université, INSERM, CNRS, Université d'Angers, CRCI2NA, Nantes, France*
[*] *these authors contributed equally to this work*

#### Contents

**Tables S1-S3**

**Figures S1-S22**

**Movies S1**

**File S1**

| **Short name** | **Long name** | **Genetic modifications** |
|---|---|---|
| WT | CP050 | $\Delta$*fhuA* |
| Rib | AP030 | *rpsB-LEI-GFPmut2//argG-LEI-mScarlet-I//*$\Delta$*fhuA* |

**Table S1: Strains used in this study.** The strains were derived from the *E. coli* K-12 wild-type strain BW25113 [1]. Rib stands for **Rib**osomes. The Rib strain carries a second reporter protein (mScarlet-I), fused to ArgG, the enzyme involved in the biosynthesis of L-arginine. This second reporter was not used in this study.

| **Identifier** | **Medium switches** | **Strain** | **#Cells** |
|---|---|---|---|
| Ref | ace 24h $\rightarrow$ glc 16h | Rib | 129 |
| | glc 16h $\rightarrow$ ace 24h | Rib | 130 |
| | ace 24h $\rightarrow$ glc 16h | Rib | 122 |
| Rep | ace 41h $\rightarrow$ glc 16h | Rib | 105 |
| | glc 16h $\rightarrow$ ace 24h | Rib | 103 |
| Bleach | ace 12h | Rib | 30 |

**Table S2: Experimental conditions of microfluidic experiments.** The following abbreviations were used: ace - acetate and glc - glucose. In the reference experiment (Ref), we distinguish three different phases (first upshift, downshift, and second upshift). The column "#Cells" mentions the total number of mother cells retained for analysis in each phase. Rep denotes an independent replicate experiments carried out in the same conditions. The experiment Bleach was used to estimate the autofluorescence and the effect of photobleaching.

| **Protein** | $k_{mat}$ **[1/min]** | $\gamma$ **[1/min]** | **Source** |
|---|---|---|---|
| GFPmut2 | $0.08 \pm 0.004$ | 0.0002 | [2] |

**Table S3: Values of the parameters in the model used for estimating resource allocation strategies.** The degradation parameter ($\gamma$) and maturation parameter ($k_{mat}$) characterize the dynamics of the fluorescent protein used in this study, according to Eqs 3-4 in the main text. The values were taken from previous work [2].

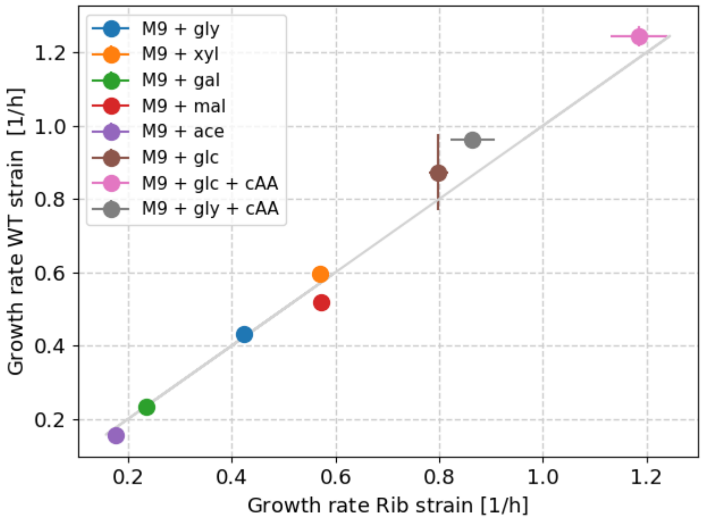


**Figure S1: Comparison of the growth rate of wild-type and reporter strains in multiple conditions.** Strain Rib was grown in a microplate reader in M9 minimal medium supplemented with several carbon sources, as detailed in the figure legend (glc - glucose, ace - acetate, gly - glycerol, fru - fructose, xyl - xylose, pyr - pyruvate, mal - maltose, gal - galactose, cAA - casamino acids). In parallel, we grew the wild-type (WT) strain (Table S1) and we monitored the absorbance of the growing cultures (*Materials and methods*). After outlier filtering and background correction, the growth rate during exponential growth was estimated from the absorbance curves using the method described by Zulkower *et al.* [3]. Each point is the mean of 5 replicates and confidence intervals are given by two times the standard error of the mean. The scatter plots show the growth rates of the wild-type strain compared with the growth rate of the two reporter strains over the different conditions (diagonal in gray). The reporter strain and the wild-type strain grow very similarly ($R^2_{Rib} = 0.99$). Therefore, no growth defect is observed after tagging the ribosomal protein RpsB with GFPmut2.

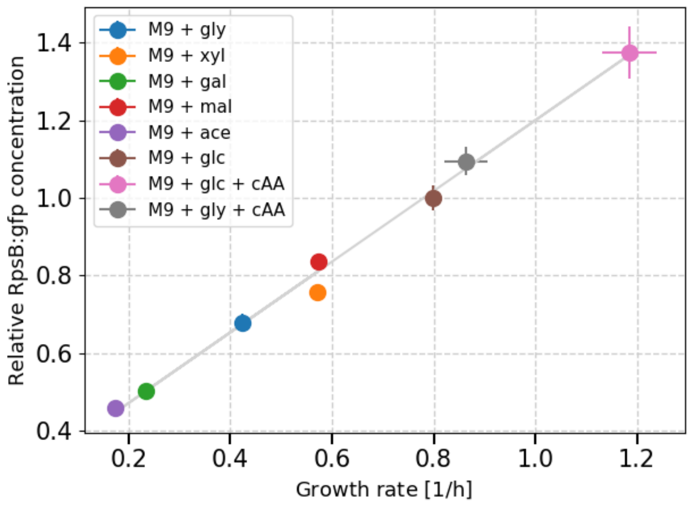


**Figure S2: Validation of growth-rate dependence of ribosome concentrations in reporter strains.** Strain Rib was grown in M9 medium supplemented with a variety of carbon sources. The same abbreviations as in Fig. S1 were used. Absorbance and green fluorescence were monitored during batch growth in a microplate reader, as described in the *Materials and methods*. After outlier filtering and background correction, the growth rates and reporter concentrations were estimated from the absorbance and fluorescence curves, using the method described by Zulkower *et al.* [4]. The reporter concentrations were then normalized by the concentration obtained in the glucose condition to obtain the relative reporter concentration. Each point is the mean of 5 replicates and confidence intervals are given by two times the standard error of the mean. Straight lines (gray) were used to fit the data and serve as visual aids for interpretation. There is a robust linear relationship between the growth rate and the RpsB-GFPmut2 concentration, as predicted by the ribosomal growth law.

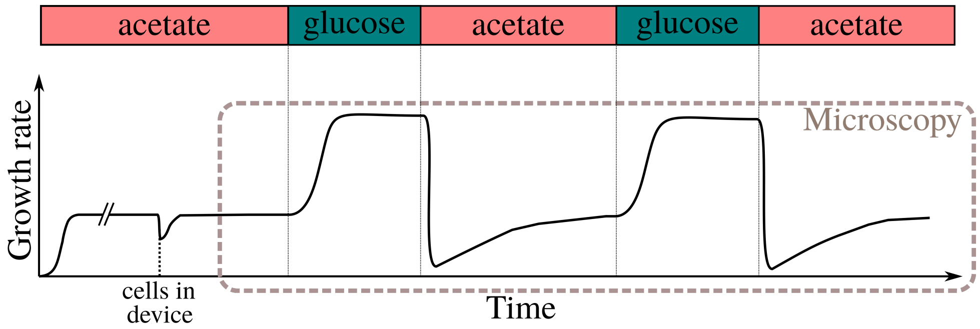


**Figure S3: Schematic outline of a typical microfluidic experiment.** Pre-cultures growing exponentially in M9 minimal medium supplemented with acetate were prepared and injected into a mother machine. A constant flow of 20 $\mu$L/min of fresh acetate medium was supplied to the growing bacteria in the device. The cell length and the green fluorescence intensity were monitored over time (*Materials and methods*). The bacteria were submitted to an upshift to glucose followed by a downshift to acetate, twice. After each shift, the bacteria were allowed to adapt to the new environment for at least eight generations. The exact scenarios of all experiments are detailed in Tab. S2.

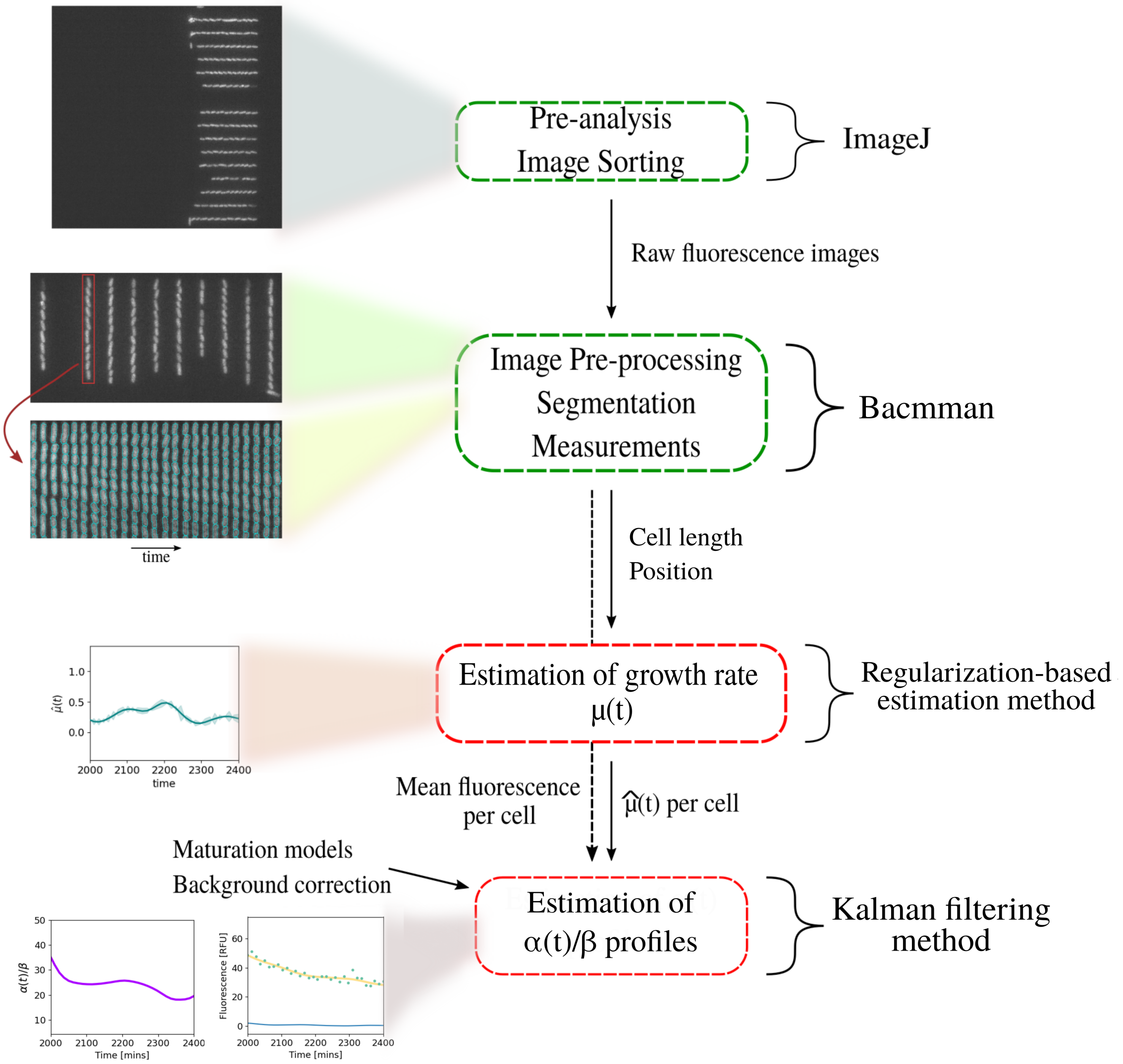


**Figure S4: Data analysis pipeline implemented for this study.** Raw fluorescence images were imported into ImageJ for image sorting. The sorted images were then imported into BACMMAN [5] for image pre-processing, and segmentation of microchannels and bacteria on the fluorescence channel. After manual checking and correction of the segmentation results, cell length and mean fluorescence intensities of all channels were determined for each cell. The cell length was used to estimate growth rate using a regularization-based fitting method in the log domain. The growth-rate estimates were used along with the background-corrected mean fluorescence intensity for each cell by a Kalman smoothing method that reconstructs the time-varying resource allocation strategies. Maturation models were used in the method to correct for fluorescent reporter dynamics. The figure shows an example image or plot for each step of the procedure. Green rounded boxes represent tools developed in previous studies, red rounded boxes represent methods developed for this study. More information on each step in the pipeline can be found in the *Materials and methods*.

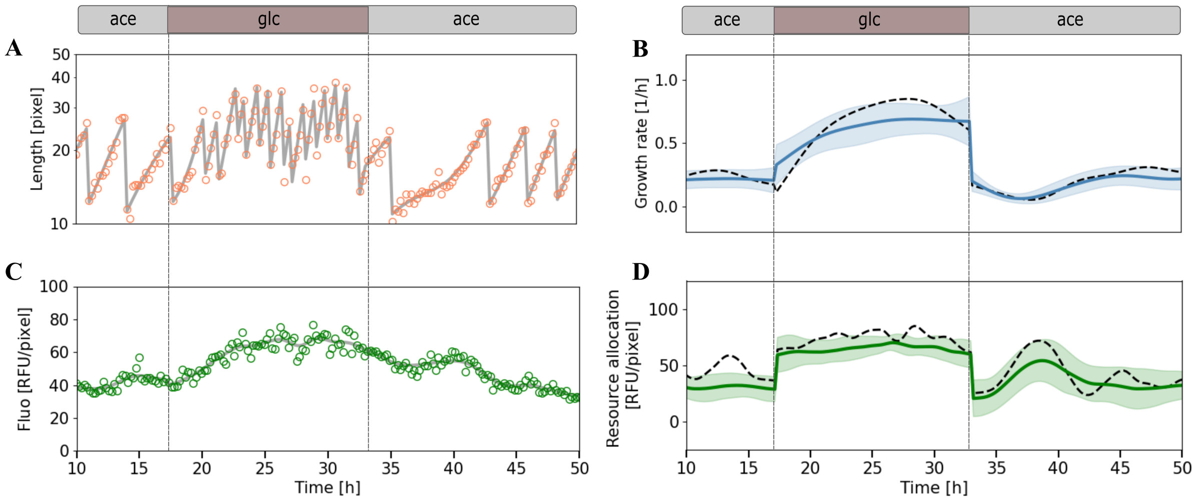


**Figure S5: Measured and estimated quantities in a replicate mother machine experiment with the ribosomal reporter strain.** The figure shows the results of a replicate experiment carried out in the same conditions as the experiment shown in Fig. 1. **A,C.** Cell length in log scale (orange dots) and green fluorescence intensity (green dots) of individual bacteria. The experiment consisted in several consecutive upshifts and downshifts (vertical dashed lines) between minimal media with glucose (glc) or acetate (ace). **B,D.** Cell length and fluorescence intensity measurements were used to estimate growth rates and resource allocation strategies, respectively, using appropriate statistical inference methods (*Materials and methods*). The gray solid curves in panels A and C represent the fits of the single-cell data obtained from the inference methods. The black dashed curves in panels B and D represent the corresponding estimates of the growth rate $\mu(t)$ and the resource allocation strategy $\alpha(t)/\beta$ for this same mother cell. Blue and green solid curves represent the mean of the estimates over all cells considered in the experiment and confidence intervals are given as two times the standard deviation. The replicate experiment reproduces the observations made for Fig. 1 in the main text.

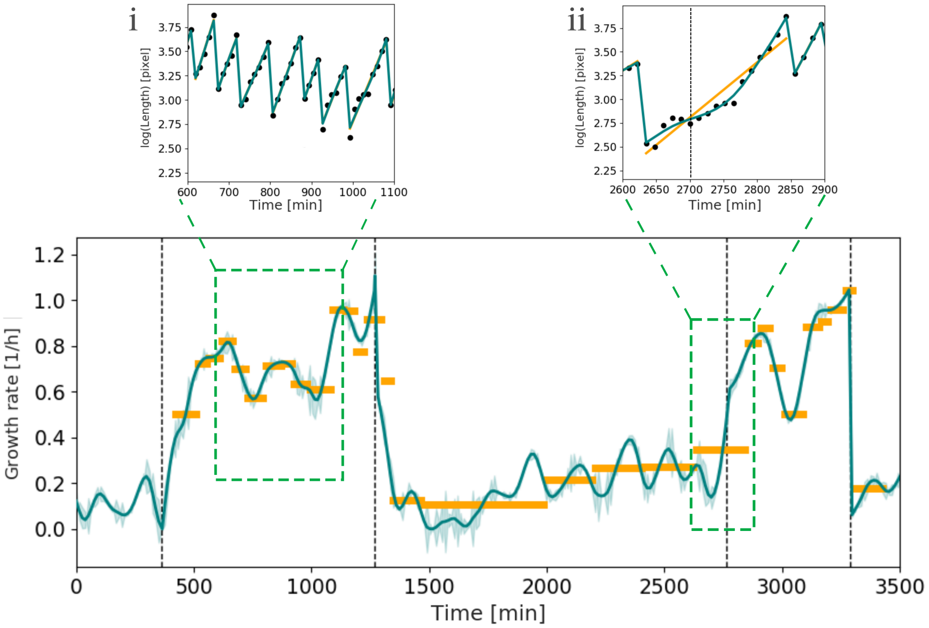


**Figure S6: Comparison of growth-rate estimation methods.** The length measurements of one mother cell were used to estimate the growth rate by means of the regularization method described in the *Materials and methods* (blue curves), for an ace-glc experiment (Tab. S2) with multiple growth transitions (dotted lines). The blue shading represents the standard errors of the estimates. This method was compared with a simple fit of an exponential curve to the measured cell lengths between divisions (orange lines) on the same data. The fits obtained from our regularization method (blue) and the exponential fit (orange) on the logarithm of length data (black dots) were compared for two distinct phases of the experiment (**i - ii**). During steady-state exponential growth, the two methods give similar estimates as shown by the superimposed fits in the inserted panel **i**. After switches between growth phases (**ii**), our method, which does not assume that growth rate is constant over a generation, produces more accurate fits, and therefore provides more reliable growth-rate estimates.

**A**

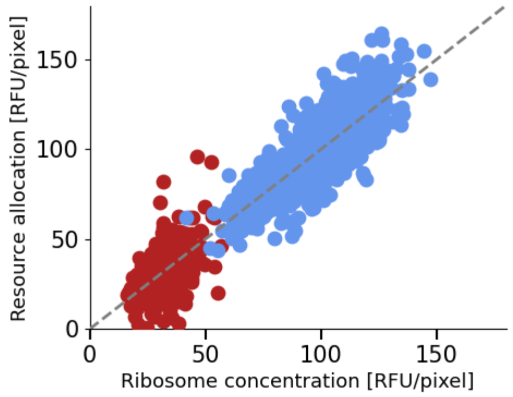


**B**

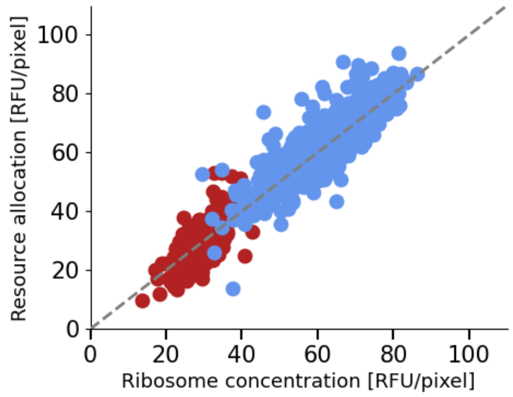


**Figure S7: Correlation between ribosome concentration and resource allocation during balanced growth. A.** Relation between (total) ribosome concentration $r$ and resource allocation strategy $\alpha/\beta$ for individual generations and individual mother cells during balanced growth of the Rib strain on acetate (red) or on glucose (blue) in the mother machine experiment shown in Fig. 1. The gray dashed line is the diagonal of the scatter plot. As expected by the model of Eq. 1, the two quantities are strongly correlated ($R^2 = 0.86$). **B.** Results for a replicate experiment with the same strain in the same growth conditions, see Fig. S5 ($R^2 = 0.88$).

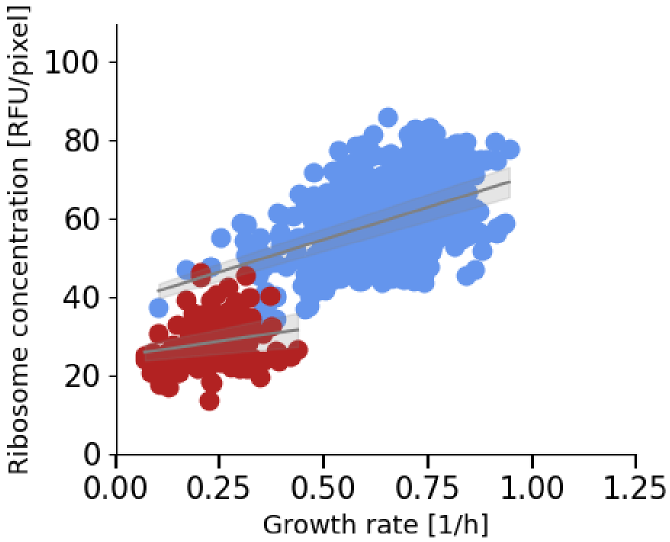


**Figure S8: Single-cell resource allocation for ribosomes during balanced growth in replicate experiment.** Relation between ribosome concentration and growth rate for individual generations of individual cells during balanced growth of the Rib strain on glucose (blue dots) and acetate (red dots) in the replicate experiment of Fig. S5. A line was fitted for the two datasets and plotted in gray, along with the confidence bands that represent two times the standard deviation. Like in the reference experiment shown in Fig. 3 in the main text, we observe a weak dependence of the ribosome concentration and growth rate on the single-cell level: $14.3 \pm 6.3$ (acetate) and $31.7 \pm 2.3$ (glucose).

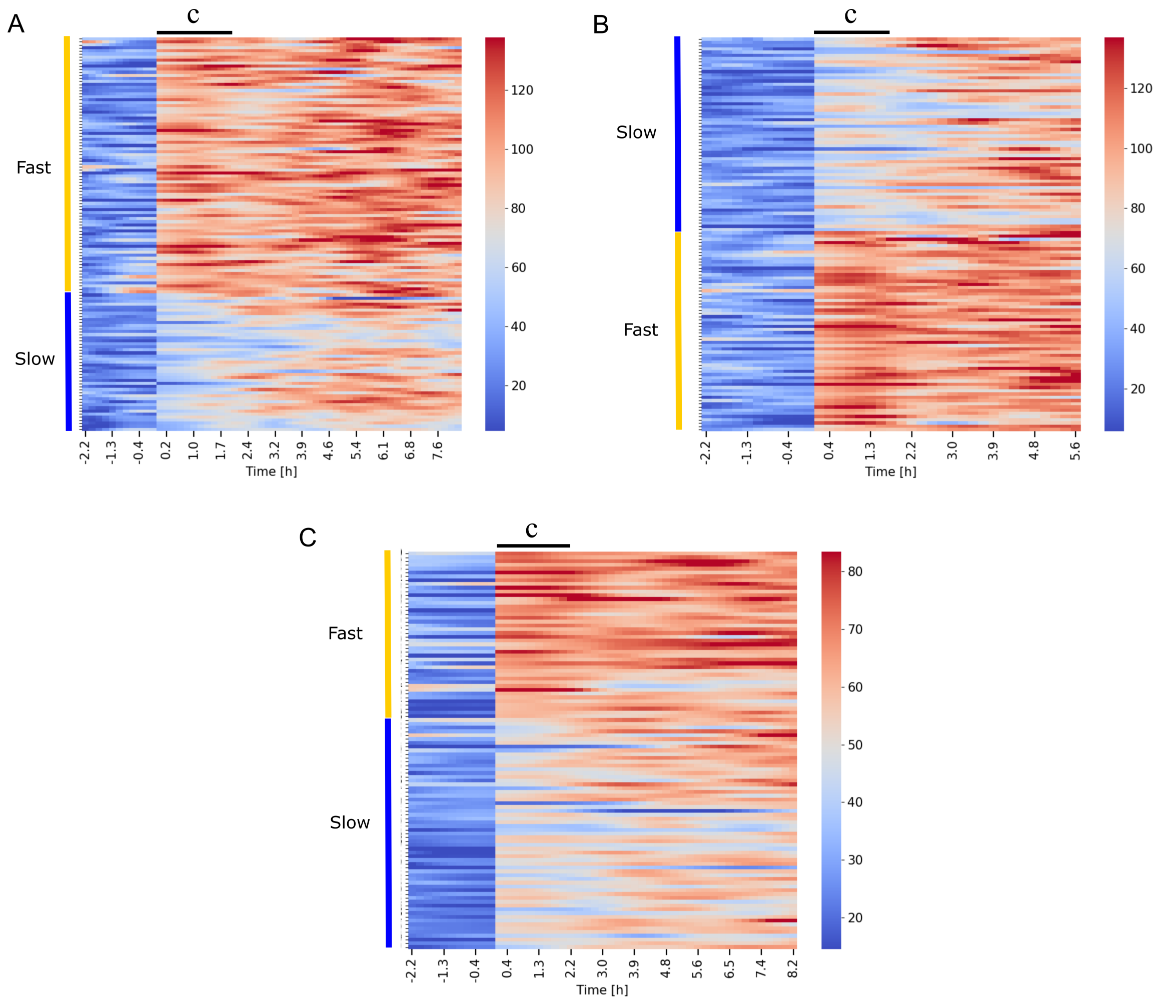


**Figure S9: Heatmaps of clustered single-cell resource allocation strategies after a nutrient upshift.** Time-varying resource allocation strategies after an acetate-glucose upshift, inferred from the fluorescence data for individual cells, were clustered by means of the k-means algorithm (*Materials and methods*). Each heatmap corresponds to an upshift in a specific mother machine experiment. Each line in a heatmap corresponds to the value of $\alpha/\beta$ (in color code, from blue to red) for a specific cell as a function of time, with time 0 indicating the time of the upshift. **A.** First upshift with the Rib strain carrying the fusion protein RpsB-GFPmut2, corresponding to the reference experiment shown in Fig. 4 of the main text. **B.** Second upshift in the same experiment (Fig. S10). **C.** Upshift using the Rib strain in an independent replicate experiment (Fig. S11). The strategies were clustered over the initial time-interval after the upshift, indicated by the black lines and letter c. Time 0 indicates the time of the upshift.

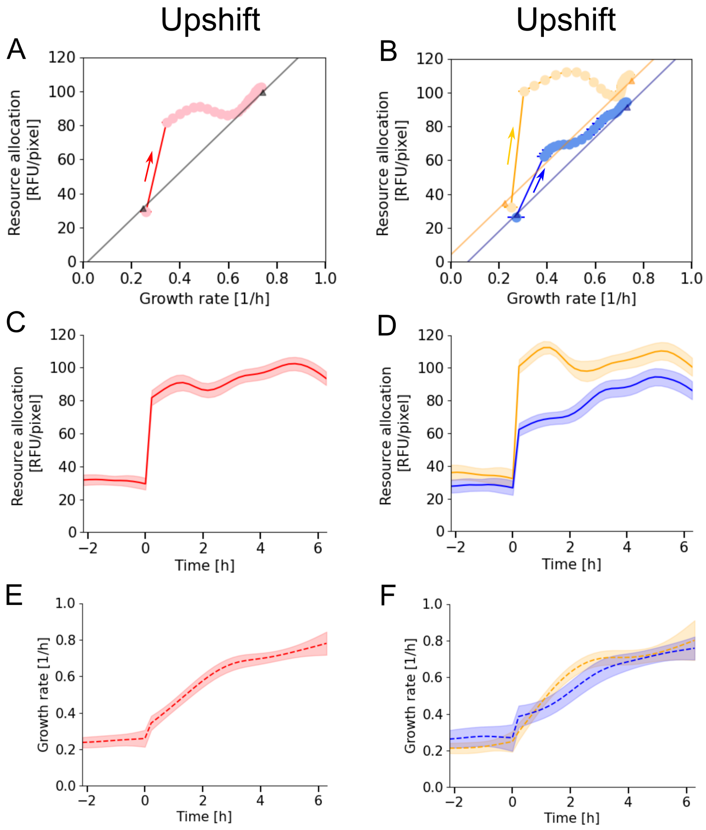


**Figure S10: Adaptation dynamics of growth rate and ribosomal resource allocation after a second nutrient upshift in the reference experiment. A.** Mean adaptation trajectory of the ribosomal resource allocation strategy $\alpha/\beta$ and the growth rate $\mu$ for a second acetate-glucose upshift applied to the Rib strain growing in a mother machine. The resource allocation strategies and growth rates were estimated from the data using the inference methods of Fig. 2 in the main text and averaged over 122 cells. The arrows indicate increasing time after the upshift. The triangles denote the average values of $\alpha/\beta$ and $\mu$ during balanced growth on acetate (before the upshift) and glucose (after the upshift). The latter values were determined by computing for each cell the mean growth rate over a period of balanced growth ($> 2$ h), and then averaging these values over the individual cells. The black line through the population average before and after the upshift is shown as a visual aid. The trajectories show that the adaptation of resource allocation and growth rate are uncoupled in a first phase and coupled in a second, like in Fig. 4 in the main text. **B.** Clustering of the single-cell resource allocation strategies for ribosomes after the upshift using k-means (*Materials and Methods*) reveals two distinct trajectories for fast adapters (orange, 62 cells) and slow adapters (blue, 60 cells). The orange and blue straight lines connect the balanced growth values (triangles) of the two populations before and after the shift. **C-D.** Time-courses of the resource allocation strategies in panels A-B, respectively. **E-F.** Time-courses of the growth rates in panels A-B.

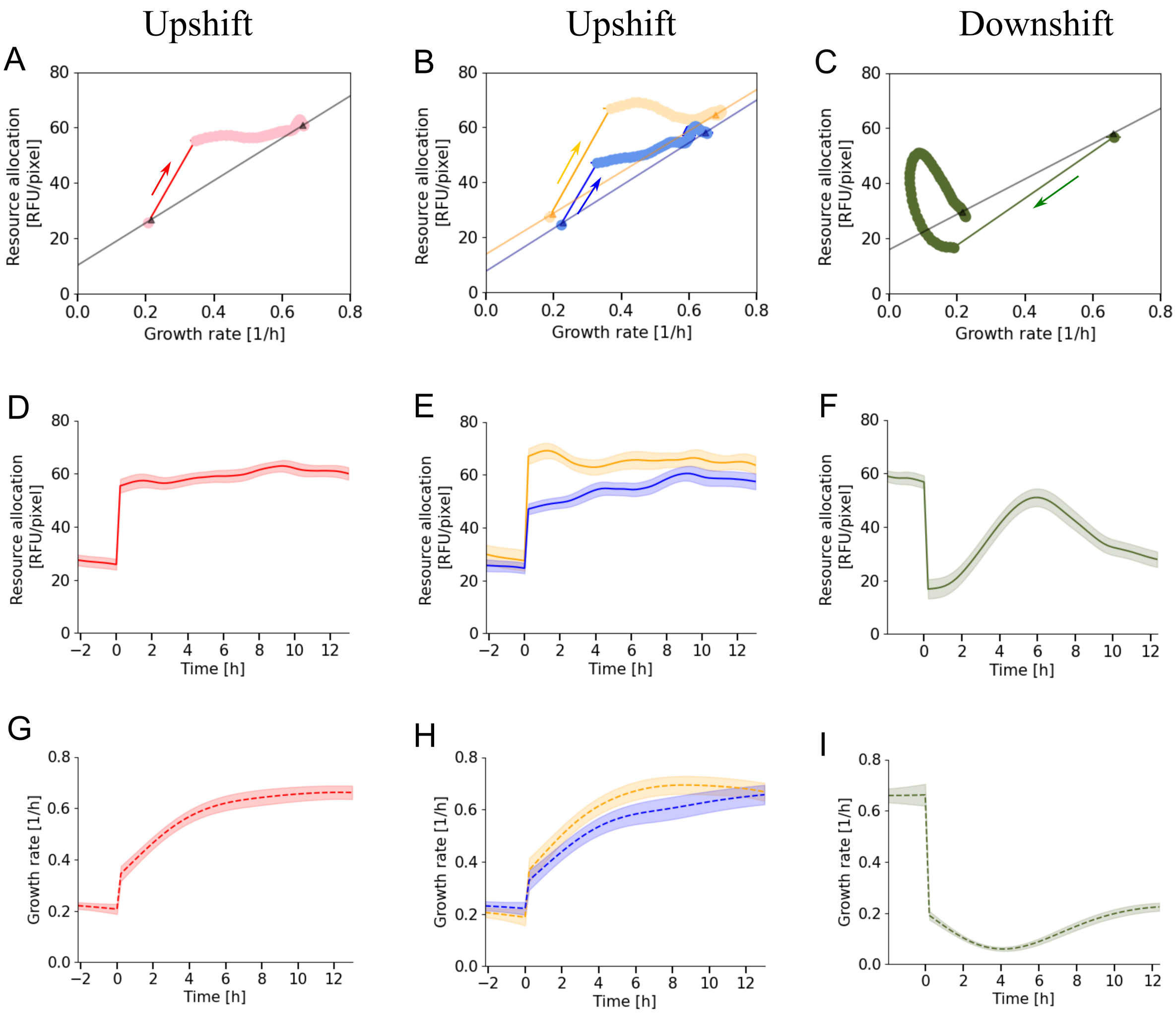


**Figure S11: Adaptation dynamics of growth rate and ribosomal resource allocation after a nutrient upshift and downshift in an independent replicate of the reference experiment.** The panels and graphical conventions are the same as in Fig. S10 (with a total of 105 cells, 61 slow adapters and 44 fast adapters). The results of this replicate experiment, using the Rib strain carrying the fusion protein RpsB-GFPmut2, agree with those observed in the reference experiment shown in Fig. 4 in the main text. The main difference lies in the time-ordering of the initial drop in growth rate and resource allocation, after the downshift to acetate. As explained in the main text, this discrepancy is probably due to the fact that the sampling density in this experiment (one measurement every 13 minutes) was too low to reliably capture events occurring in such rapid succession.

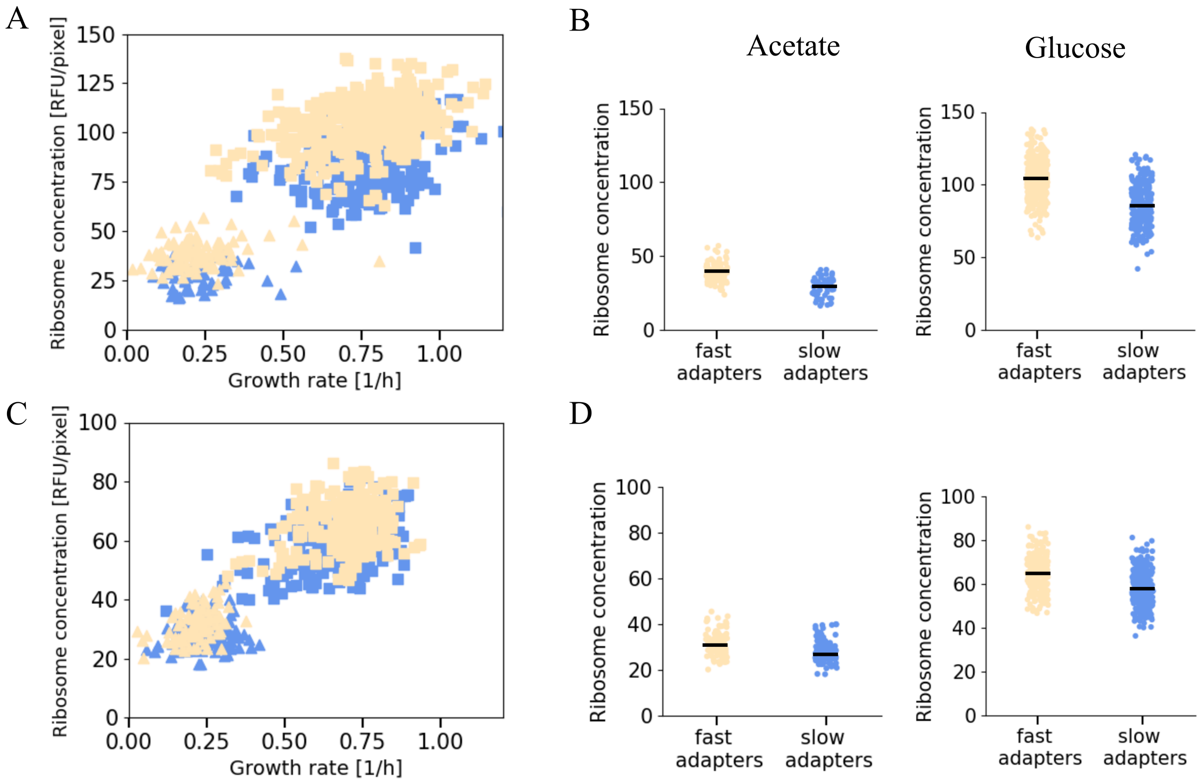


**Figure S12: Comparison of ribosome concentrations in fast and slow adapters before nutrient upshifts. A.** Relation between growth rates and ribosome concentrations for individual generations of individual cells of the Rib strain during balanced growth on acetate or glucose (triangles and squares, respectively). The data points corresponding to fast and slow adapters in the two generations before the upshift from acetate to glucose are colored in orange and blue, respectively. Idem for the data points corresponding to fast and slow adapters after 6 generations following the upshift. **B.** Violin plots of ribosome concentrations in panel A, for fast and slow adapters. The non-parametric Mann-Whitney U test shows that the violin plots correspond to non-identical distributions, with the distribution of fast adapters stochastically higher than the distribution of slow adapters (p-values: $1.9 \cdot 10^{-15}$ and $2.9 \cdot 10^{-47}$ for growth on acetate and glucose, respectively). In particular, the median ribosome concentrations are higher in fast adapters than in slow adapters, suggesting the availability of a higher ribosome reserve before the upshift. **C-D.** Idem for a replicate experiment with the same strain (data from Fig. S8A, p-values: $6.2 \cdot 10^{-6}$ and $1.3 \cdot 10^{-15}$ for growth on acetate and glucose, respectively).

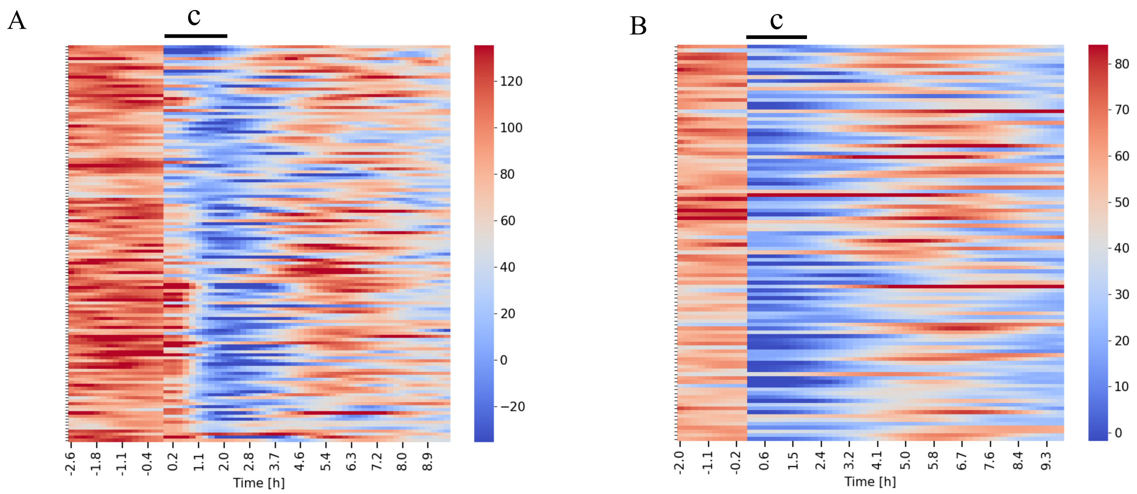


**Figure S13: Heatmaps of single-cell resource allocation strategies after a nutrient downshift.** Time-varying resource allocation strategies after an acetate-glucose downshift with the Rib strain were inferred from the fluorescence data for individual cells, obtained in different mother machine experiments. **A.** Downshift in the reference experiment, as shown in Fig. 4 of the main text. **B.** Downshift in an independent replicate experiment (Fig. S11). Clustering over the initial time-interval after the upshift, indicated by the black lines and letter c, did reveal some heterogeneity in the case of the reference experiment, but not in the replicate experiment where the temporal resolution of the measurements directly after the downshift was lower.

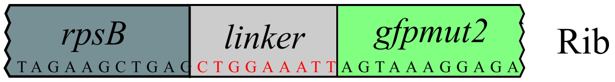


**Figure S14: Reporter strain used in this study.** The reporter strain was constructed using lambda Red-mediated homologous recombination via the intermediate of a selection "cassette", as described in the *Materials and methods*. The reporter gene (*gfpmut2* was fused with the target gene *rpsB* by means of a flexible linker of three amino acids. The complete genotype of the strain is described in Tab. S1.

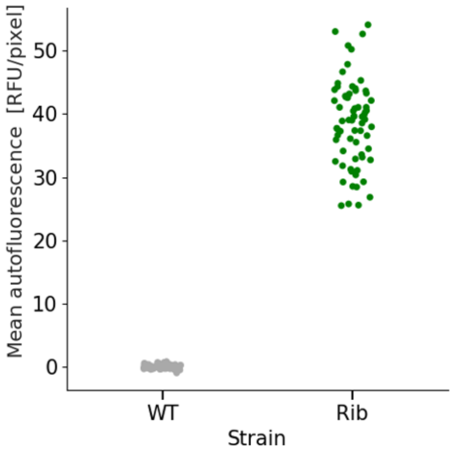


**Figure S15: Quantification of the autofluorescence of the reporter strain and the wild-type strain.** Strains Rib and WT were inoculated on an agar pad containing M9 minimal medium supplemented with 2 g/L of acetate and left to grow overnight, as described in the *Materials and methods*. Images of green fluorescence were acquired for the Rib and WT strains in parallel. Image acquisition parameters were used as described in the *Materials and method*. For each strain, 70 cells were segmented using ImageJ and the mean fluorescence intensity [RFU/pixel] was determined for each cell over a sequence of frames. For each image, the background was removed by subtracting the fluorescence intensity of the pad background. The mean fluorescence intensity of the WT strain is 100-4000-fold lower than the fluorescence of the modified strains, and therefore negligible for the purpose of our study.

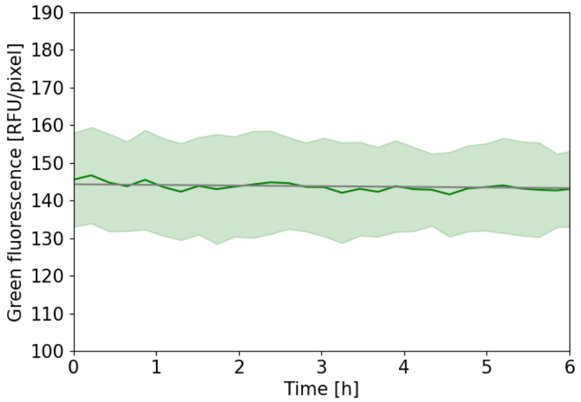


**Figure S16: Quantification of the effect of photobleaching for the fluorescent reporter used in this study.** A microfluidics control experiments with the Rib strain was conducted in M9 minimal medium supplemented with 2 g/L of acetate. The experimental protocol, image acquisition parameters and segmentation procedure are described in the *Materials and methods*. The graphs present the mean of green fluorescence intensities for 30 mother cells along with 2 times the standard error of the mean. Given that the strains are in balanced growth, with a constant average reporter concentration, we expect any bias due to photobleaching to reveal itself by a decrease of the fluorescence intensity. A straight line was used to fit the mean curve in Python. The slope obtained (in unit RFU/(pixel h)) was $-0.16 \pm 0.47$. The coefficient is indistinguishable from zero, so no photobleaching correction was applied to the data.

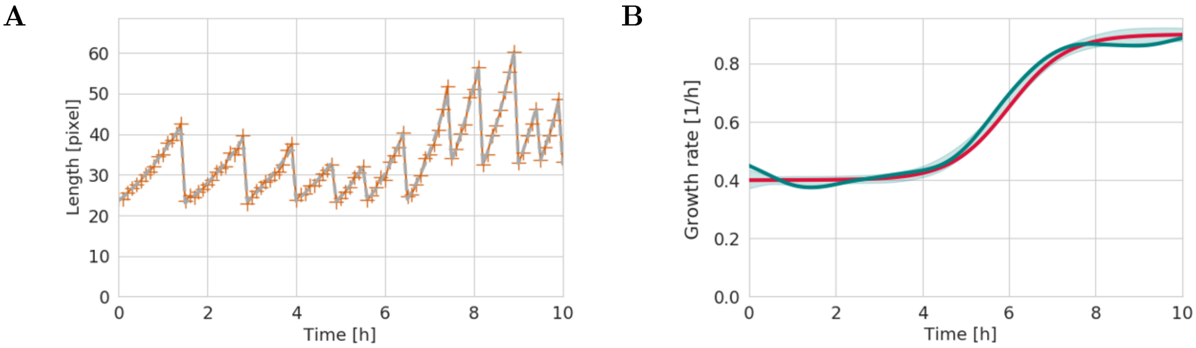


**Figure S17: Performance of growth-rate estimation method on synthetic data.** A sigmoid-like function was defined to resemble a transition from low to high growth rate. A series of cell division points, along with cell lengths after division with some added white noise were defined to resemble a transition from a poor to a rich carbon source, as observed in our microfluidics experiments. **A.** Using the model for cellular growth described in the *Materials and methods* of the main text (Eq. 5), and the defined cell division points, a series of length measurements was generated with added white noise ∼N(0, 1.4) (orange points). **B.** The growth-rate estimation method described in the *Materials and methods* and Text S2 was used to provide regularized estimates of the growth rate. The corresponding fit to the length data is shown in panel A (gray curve) and the resulting growth rate curve in panel B (blue curve). The shaded blue area represents two times the standard error of the mean of growth-rate estimates obtained from 100 generated datasets. The method is shown to provide growth-rate estimates that overlap the true growth rate (red curve) used as input for the simulations.

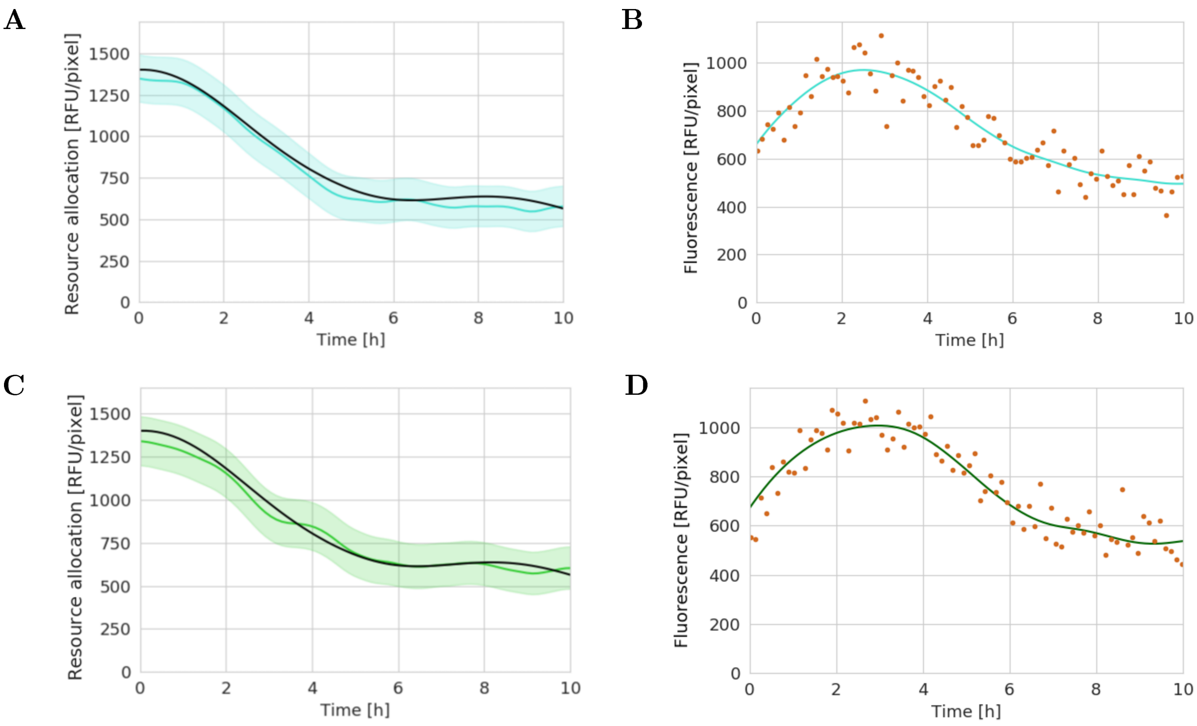


**Figure S18: Performance of Kalman smoothing algorithm on synthetic data.** (**A**) A sinus-like promoter activity $pa(t) = (\mu(t) + \gamma) \cdot \alpha(t)/\beta$ was generated and divided by the sum of the simulated growth rate (Fig. S17B, red curve) and degradation constant $\gamma$ to obtain the simulated resource allocation strategy (black curve). This curve was compared with the estimated resource allocation profile (blue curve), obtained by applying the Kalman smoothing method (*Materials and methods* and Text S2) to the simulated data. The shaded blue area represents the confidence interval of the estimates produced by the method (Text S2). (**B**) Synthetic fluorescence intensity data generated by means of Eqs 3-4 in the main text and the resource allocation profile in panel A, with added white noise of amplitude equivalent to that observed in the single-cell fluorescence measurements (∼N(0, 80), orange points). The blue curve is the fluorescence intensity predicted from the estimated resource allocation strategy. As can be seen in panels A and B, the algorithm is able to robustly reconstruct the resource allocation profile from the synthetic data. (**C** - **D**) To verify that the method is able to reconstruct the resource allocation strategy using estimates of the growth rate, $\hat{\mu}(t)$ rather than the true growth rate $\mu(t)$, the resource allocation profile was reconstructed using synthetic data generated from the same sinus-like promoter activity as above, but divided by the growth rate estimates $\hat{\mu}(t)$ of Fig. S17B (blue curve). The use of growth-rate estimates instead of the true growth rate does not affect the quality of the reconstruction of the resource allocation strategies and the correspondence of the predicted and observed fluorescence intensities.

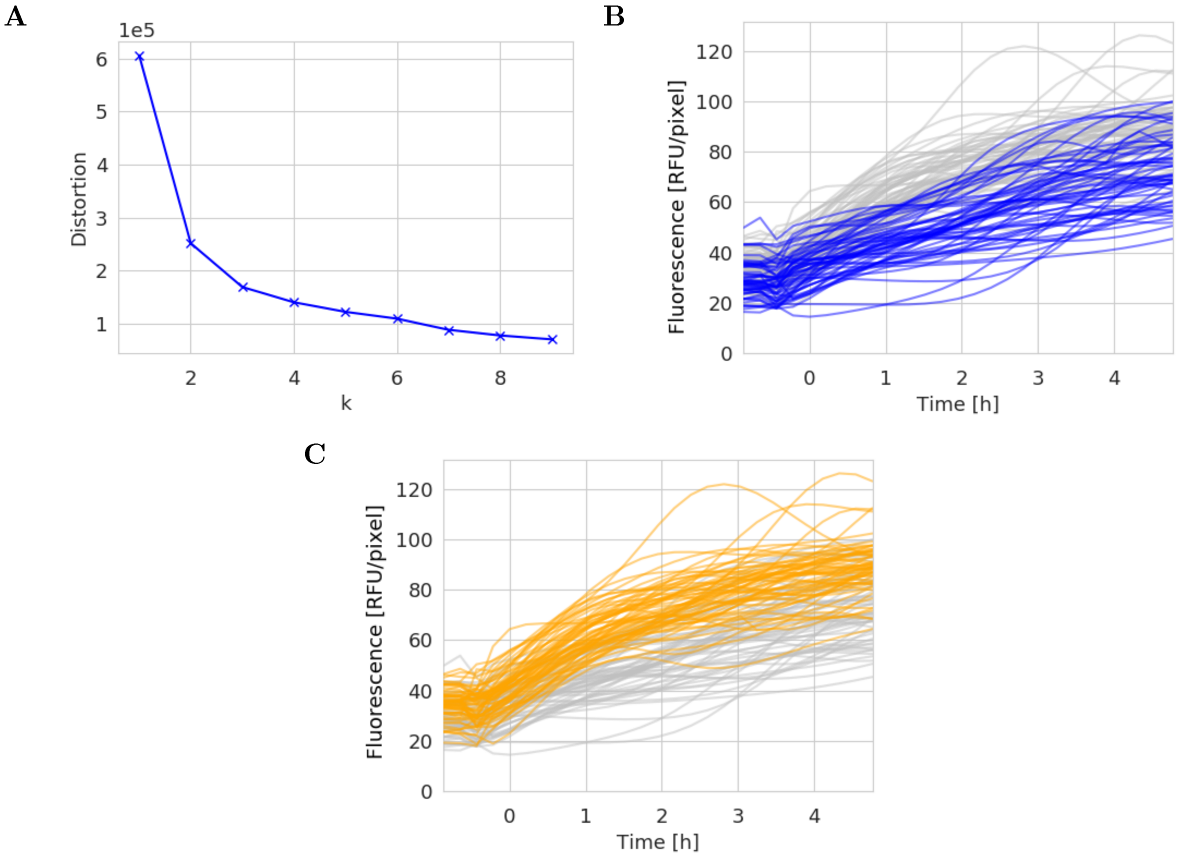


**Figure S19: Determination of the number of clusters used by the k-means algorithm.** In order to evaluate the variability of the adaptation dynamics of ribosomes, we applied the k-means clustering algorithm to the inferred resource allocation strategies after an acetate-glucose upshift (*Materials and methods* and Fig. S9). We decided on the number of clusters ($k$) appropriate for the data by relying on the so-called elbow method, often used in the context of k-means clustering [6]. As a metric for the information added by each cluster, we used cluster distortions [7]. The optimal number of clusters is given by the "elbow" in the curve displaying the distortion as a function of $k$, where adding another cluster significantly decreases the marginal gain in information. In our case, the optimal number of clusters is 2 or 3 (**A**). The two clusters correspond to two clearly distinguishable patterns in the fluorescence intensities (**B-C**).

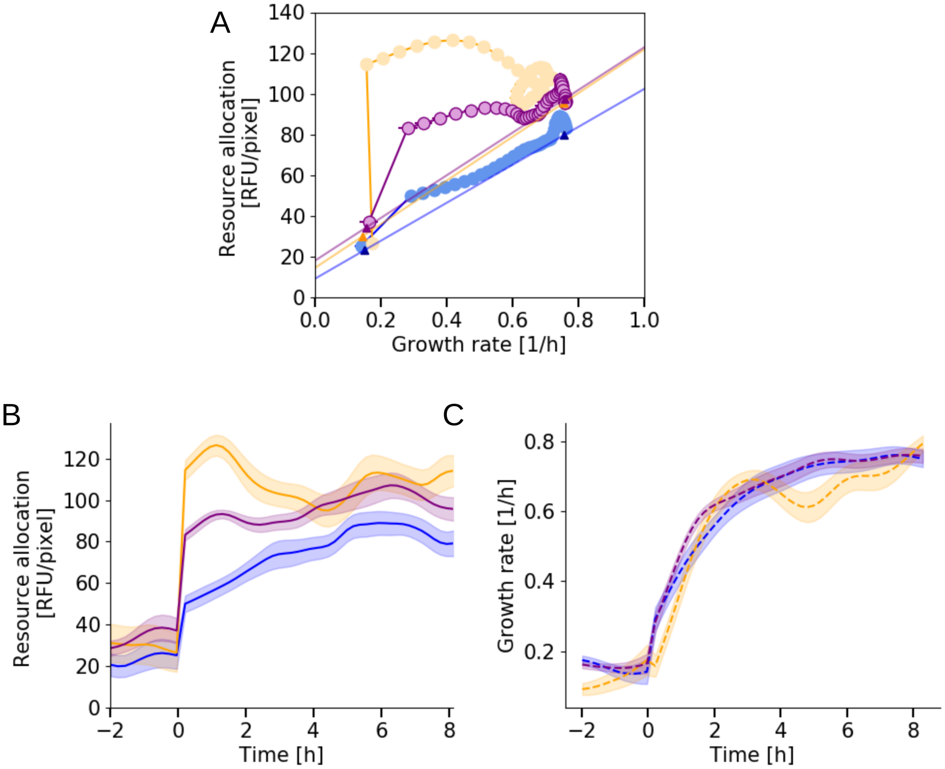


**Figure S20: Adaptation dynamics of growth rates and ribosomal resource allocation strategies in the case of three clusters A.** Adaptation trajectory of ribosomes after an acetate-glucose upshift for the data presented in Fig. 4 of the main text but for three instead of two clusters. The k-means clustering algorithm was used, as described in the *Materials and methods*. **B-C.** Corresponding time-courses of the mean resource allocation strategies (panel B) and mean growth rates (panel C) for the three clusters. Confidence intervals are given by two times the standard error of the mean. The yellow and purple clusters have the same resource allocation for balanced growth on glucose or acetate, and the same growth rate during the upshift. The distinction of three rather than two clusters therefore does not add any useful information from a biological point of view.

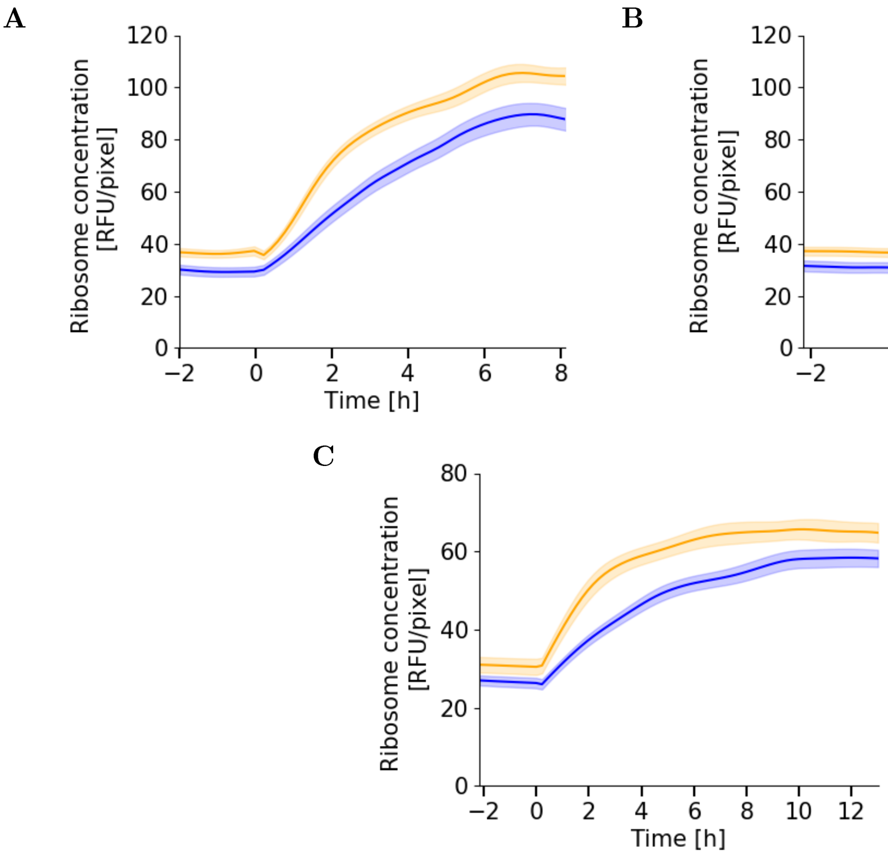


**Figure S21: Adaptation dynamics of ribosome concentration after nutrient upshift for slow and fast adapters. A.** Time-courses of the mean total ribosome concentrations of fast (orange curve) and slow (blue curve) adapters after the acetate-glucose upshift for the Rib strain growing in a mother machine in the reference experiment. The corresponding resource allocation strategies are shown in Fig. 4E in the main text. The total ribosome concentrations were estimated from the data using the inference methods of Fig. 2. **B.** Idem for a second upshift in the same experiment (resource allocation strategies in Fig. S10D). **C.** Idem for an independent replicate of the reference experiment (resource allocation strategies in Fig. S11E). In all experiments the fast adapters maintain a higher ribosome concentration after the upshift, in agreement with the higher proportion of resources allocated to ribosomal synthesis (Text S3).

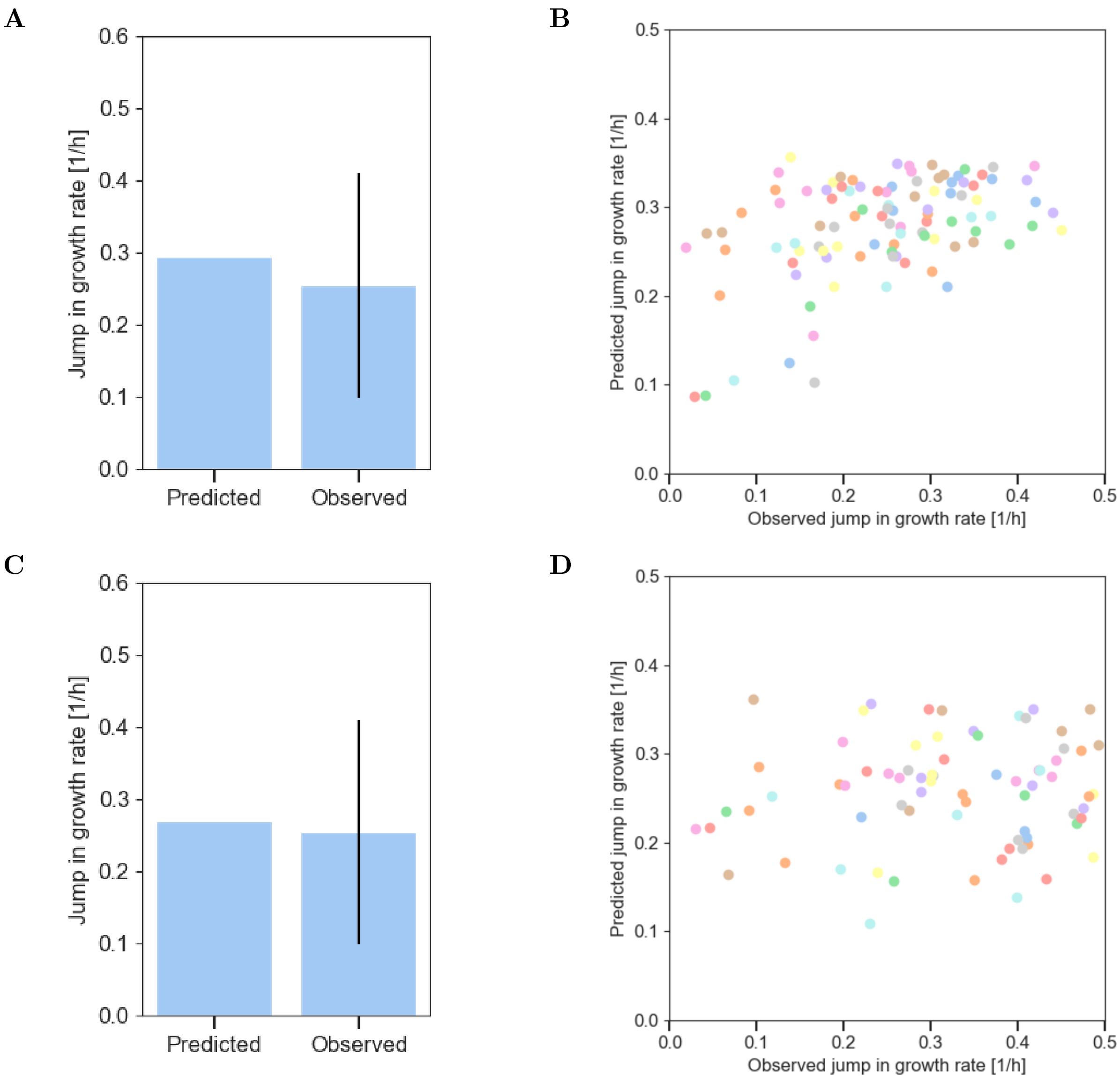


**Figure S22: Predicted and observed jump in growth rate after nutrient upshift.** We experimentally tested the predictions of the jump in growth rate directly after the glucose-acetate upshift in the reference experiment, using the theoretical models of Korem Kohanin *et al.* [8] and Mori *et al.* [9], as explained in detail in the *Materials and methods* section. **A.** We applied the model of Korem Kohanin *et al.* to population-averaged values of the growth rate directly before the upshift and the growth rate during balanced growth on glucose after the upshift. The predicted population-averaged value of the growth rate directly after the upshift ($\mu_1$) was compared with the observed value, computed from the experimental data. The uncertainty interval for the experimentally determined value is given by the standard deviation. There is a good correspondence between the predicted and observed values on the population level. **B.** Idem, a scatter plot comparing the values for the individual cells. There is no correspondence between the predicted and observed values on the single-cell level. **C-D.** Like in panels A-B, but for the model Mori *et al.*

**Movies S1: Time-lapse phase contrast and fluorescence images of nutrient upshift and downshift.** The two movies (`movie_phase.avi` and `movie_fluo.avi`) show the glucose-acetate upshift in the reference experiment, using the Rib reporter strain. The times shown in the movies correspond to the time axes in Fig. 1 in the main text.

**File S1: Python code for estimating growth rates and resource allocation strategies, as well as for generating the figures in the main text and the supplementary information.** The Python estimation code consists of two files (`estimation.py` and `smoothing.py`). `estimation.py` reads the files with time, length and fluorescence data for each mother cell (in the directory .\`REF/REP_data_mothercells`), the file with time-points at which a cell division occurs (`REF/REP_divisions.csv`), and the file with experimental parameters (`REF/REP_params.csv`). The code performs the growth-rate and resource allocation estimation procedures outlined in Text S2, calling `smoothing.py` for the automatic tuning of regularization parameters. The output of `estimation.py` is a directory of estimation results .\`REF/REP_estimates_output`, containing for each mother cell the estimated values of growth rate and resource allocation over time, as well as other statistics. The file `analysis.py` takes the latter file as input to produce the figures in the main text and the supplementary information. Both `estimation.py` and `analysis.py` are parametrized with the identifier of the experiment considered (`REF` or `REP`).
