## Supplementary material for "Single-cell data reveal heterogeneity of resource allocation across a bacterial population": SI texts

### Supplementary Texts for "Single-cell data reveal heterogeneous resource allocation across a bacterial population"

### Contents

**Text S1 Model definition** **2**

**Text S2 Estimation of growth rates and resource allocation strategies** **5**

**Text S3 Model analysis** **14**

### Text S1 Model definition

This supplementary text describes how the dynamical models used for the inference of ribosomal resource allocation strategies in the main text can be derived from basic biological assumptions. We proceed in two steps. In the first step, we relate resource allocation strategies to the dynamics of the total concentration of ribosomes. In the second step, we couple this biological model with a measurement model, in order to relate the resource allocation strategies to the observed fluorescence in single-cell time-lapse microscopy experiments. While we develop the models for the case of ribosomes, the same reasoning applies to other proteins.

#### Resource allocation model

Let $\mathrm{Vol}(t)$ [L] be the volume of the growing bacterial population at time $t$ [h]. We assume that the volume is proportional to the biomass. That is, the biomass density is constant, an assumption justified by steady-state data over a large range of growth rates in *E. coli* [1]. Moreover, we approximate the biomass by the protein mass, its major component [1, 2]. Accordingly, we define

$$\mathrm{Vol}(t) = \beta\,(R(t) + P(t)), \tag{S1.1}$$

where $R$ [g] is the total mass of ribosomes in the growing bacterial population and $P$ [g] the total mass of other proteins. $1/\beta$ [g/L] is the biomass density or, equivalently in this case, the total protein concentration.

Let $V_{prot}(t)$ be the total protein synthesis rate in the growing bacterial population, with units g/h. We now introduce a resource allocation strategy $\alpha(t)$ that attributes a time-varying fraction (between 0 and 1) of $V_{prot}$ to the synthesis of ribosomal proteins. By definition, $1 - \alpha(t)$ is the fraction of resources allocated to the other proteins.

The following system of differential equations describes the dynamics of the protein masses:

$$\frac{d}{dt}R(t) = \alpha(t)\,V_{prot}(t) - \gamma\,R(t), \tag{S1.2}$$

$$\frac{d}{dt}P(t) = (1 - \alpha(t))\,V_{prot}(t) - \gamma\,P(t), \tag{S1.3}$$

where $\gamma$ [1/h] is the protein degradation constant. Half-lives of proteins are generally long (>10 h) [3, 4]. We complete the model with the dynamics of the volume of the growing bacterial population:

$$\frac{d}{dt}\mathrm{Vol}(t) = \mu(t)\,\mathrm{Vol}(t). \tag{S1.4}$$

The variables in the above model are extensive quantities, computed over the entire growing population. For our purpose, it is more convenient to define models based on intensive quantities, that is, to normalize the protein masses by the population volume. Accordingly, we define protein concentrations $r = R/\mathrm{Vol}$ and $p = P/\mathrm{Vol}$, with units g/L, and the specific protein synthesis rate $v_{prot} = V_{prot}/\mathrm{Vol}$, with unit g/(L h).

The above definitions allow, first of all, to rewrite the growth rate as the net (specific) protein synthesis rate. Using the definition of the growth rate (Eq. S1.4) and the proportionality of biomass and volume (Eq. S1.1), we find that

$$\begin{aligned}\mu(t) &= \frac{\beta\left(\frac{d}{dt}R(t) + \frac{d}{dt}P(t)\right)}{\mathrm{Vol}(t)} = \beta\left(\frac{V_{prot}(t)}{\mathrm{Vol}(t)} - \gamma\frac{R(t)+P(t)}{\mathrm{Vol}(t)}\right) \\ &= \beta\, v_{prot}(t) - \gamma. \end{aligned} \tag{S1.5}$$

That is, the growth rate equals the total protein synthesis rate relative to the total protein concentration $1/\beta$, minus the protein degradation rate.

Second, using Eqs S1.2-S1.3, as well as the definition of the growth rate, the dynamics of the protein concentrations can be written as follows:

$$\begin{aligned}\frac{d}{dt}r(t) &= \frac{\frac{d}{dt}R(t)\,\mathrm{Vol}(t) - R(t)\,\frac{d}{dt}\mathrm{Vol}(t)}{\mathrm{Vol}(t)^2} = \frac{\frac{d}{dt}R(t)}{\mathrm{Vol}(t)} - \mu(t)\,\frac{R(t)}{\mathrm{Vol}(t)} \\ &= \alpha(t)\,v_{prot}(t) - (\mu(t)+\gamma)\,r(t), \end{aligned} \tag{S1.6}$$

$$\begin{aligned}\frac{d}{dt}p(t) &= \frac{\frac{d}{dt}P(t)\,\mathrm{Vol}(t) - P(t)\,\frac{d}{dt}\mathrm{Vol}(t)}{\mathrm{Vol}(t)^2} = \frac{\frac{d}{dt}P(t)}{\mathrm{Vol}(t)} - \mu(t)\,\frac{P(t)}{\mathrm{Vol}(t)} \\ &= (1-\alpha(t))\,v_{prot}(t) - (\mu(t)+\gamma)\,p(t). \end{aligned} \tag{S1.7}$$

Note that when defining the protein dynamics on the concentration level, protein synthesis is balanced by protein degradation and a term accounting for growth dilution [5].

The model can be further simplified by taking into account the proportionality of the protein synthesis rate with the sum of the growth and degradation rates (Eq. S1.5), which gives the following equation for the ribosomal protein dynamics:

$$\frac{d}{dt}r(t) = (\mu(t)+\gamma)\left(\frac{\alpha(t)}{\beta} - r(t)\right). \tag{S1.8}$$

The interest of this model lies in that it relates the ribosomal resource allocation strategy to the ribosome concentrations. In particular, when knowing the degradation rate and observing the time-varying growth rate and ribosomal protein concentration, we can estimate $\alpha/\beta$. In general, no reliable value for $\beta$ will be available, so that we cannot compute an absolute value for the resource allocation strategy $\alpha(t)$. When we are only interested in changes in $\alpha(t)$, however, an estimate of the normalized strategy $\alpha/\beta$ is sufficient.

#### Measurement model for the inference of resource allocation strategies

The observation of the ribosome concentration is indirect, involving the fusion of a fluorescent reporter protein with a representative ribosomal subunit, RpsB. The reporter protein has its own dynamics, in particular due to maturation of the fluorophore and photobleaching [6], which requires the resource allocation model of Eq. S1.8 to be coupled with a dedicated measurement model. In this section, we extend the model to account for maturation effects. The effects of photobleaching were found to be negligible in our conditions (*Materials and methods* and Fig. S16), so we ignored these in the analysis. While we focus on the case of fluorescent tags of ribosomes, the approach is generic and applies to other protein categories as well. The measurement models are based on previous work on the maturation of fluorescent proteins [6, 7].

In the case of green fluorescent protein (GFP), the maturation dynamics adds a second equation to the model:

$$\frac{d}{dt} r_{im}(t) = (\mu(t) + \gamma) \left( \frac{\alpha(t)}{\beta} - r_{im}(t) \right) - k_{mat}\, r_{im}(t), \tag{S1.9}$$

$$\frac{d}{dt} r_{m}(t) = k_{mat}\, r_{im}(t) - (\mu(t) + \gamma)\, r_{m}(t), \tag{S1.10}$$

where $r_{im}$ and $r_m$ refer to the concentrations of ribosomes tagged with immature and mature GFP, respectively [g/L]. The total ribosome concentration is given by the sum of the concentrations of mature and immature fusion proteins ($r_{im} + r_m$). Maturation is modeled as a first-order reaction with a constant $k_{mat}$ [1/h]. The stability of ribosomes with both the mature and the immature fluorescent tags is assumed to be the same, as determined by the degradation constant $\gamma$ [1/h].

The units of the ribosome concentrations in the above model are taken to be grams per liter. The observations of fluorescent proteins in the fluorescence microscopy images are in different units: namely Relative Fluorescence Units (RFU) per pixel (*Materials and methods*). We make the assumption that the two scales are linearly related. Multiplying the concentration variables $r_{im}$ and $r_m$, as well as the total protein concentration $1/\beta$, by a scaling factor $\theta$ [RFU/pixel $\cdot$ L/g] changes the concentration units, but not the structure of the model of Eqs S1.9-S1.10. In order to avoid unnecessarily complex notation, we keep the same variable names in the models with the original and rescaled units.

As explained in the *Materials and methods*, we have carried out targeted calibration experiments in previous work [7] to determine the values of the maturation and degradation constants of the fluorescent reporter used in this study. The results are shown in Tab. S3. The model of Eqs S1.9-S1.10 was used for the inference of the resource allocation strategies, as described in the *Materials and methods* and Text S2.

### Text S2 Estimation of growth rates and resource allocation strategies

This supplementary text describes the methods developed for the inference of a time-varying resource allocation profile from single-cell time-course measurements of fluorescent reporter intensities and cell lengths. The method relies on the models presented in the *Materials and methods* and Text S1, where the dynamics of reporter abundance $r_m(t)$ depends on the unknown resource allocation profile $\alpha(t)$ and the growth rate $\mu(t)$. Our approach relies on estimating $\mu(t)$ from cell-length data first, and then using this estimate in Eqs 3-4 in the main text to reconstruct the profile of $\alpha$ from fluorescence data. Our methods apply to so-called mother cells sitting at the dead-end of a mother machine channel and observed over a large number of consecutive generations. Before describing the estimation methods, in a first section we discuss the relation between the single-cell measurements and the quantities $r_m(t)$ and $\mu(t)$, and establish some notation. Leveraging regularization [8, 9], the estimation methods are discussed in the next two sections, with subsections discussing the automated choice of regularization parameters and the application of the methods to data from upshift and downshift experiments (Tab. S2).

#### Relation between single-cell models and experimental observations

After image processing (*Materials and methods*), we have a series of length measurements (a sawtooth-type profile because of cell growth and divisions, see Fig. 1B in the main text) along with the mean fluorescence intensities over the observed cell area. For $i = 1, \ldots, m$, let $t_{i,1}, t_{i,2}, \ldots, t_{i,n_i}$ be the $n_i$ measurement times in-between two subsequent cell divisions, with $t_{i,j} < t_{i,j+1}$ and $t_{i,n_i} < t_{i+1,1}$ for all relevant $i$ and $j$. This partitioning (corresponding to $m-1$ divisions) was obtained from the cell length measurement sequence by the segmentation procedure of BACMMAN (*Materials and methods*). We assume that $n_i \geq 2$ for all $i$, that is, that we have at least two length measurements per generation. We denote with $n = n_1 + \ldots + n_m$ the total number of measurement times for one mother cell. Correspondingly, let $L_{i,j}$ and $f_{i,j}$ be the cell length and the mean fluorescence intensity of the cell, respectively, as at time $t_{i,j}$, obtained from post-processing the image analysis results (*Materials and methods*). To relate these measurements to the model quantities $\mu(t)$ and $r_m(t)$, we make the following assumptions:

1. $L_{i,j}$ is a noisy observation of the actual cell length $L(t)$ at time $t = t_{i,j}$, where, in-between divisions, the dynamics of $L(t)$ relate to growth rate by the model

$$\frac{d}{dt}L(t) = \mu(t)\, L(t). \tag{S2.11}$$

2. The mean intensity $f_{i,j}$ is a noisy measurement of the reporter abundance per unit cell volume $r_m(t)$ at time $t = t_{i,j}$.

Note that no claim is made in Assumption 1 concerning the cell length before ($L_{i,n_i}$) and after ($L_{i+1,1}$) a division. Besides possible deviations from a symmetric division, this is because the precise division time is generally not known (it occurs between two measurement times). Also note that Assumption 2 is (up to a scaling factor) in accordance with the interpretation of $r_m(t)$ as an intensive variable, that is, the reporter concentration within the given (growing) cell at time $t$ (Text S1). Further mathematical characterization of Assumptions 1–2 is made in the appropriate sections below.

#### Growth rate estimation from cell-length data

The method developed below (see Fig. S2.1 as a reference) works on the logarithm of cell length. That is, we consider the data $\tilde{\ell}_{i,j} = \log L_{i,j}$, which are noisy measurements of $\ell(t) = \log L(t)$. For a constant growth rate $\mu$, in-between divisions, $\ell(t)$ sits on a straight line. We instead let $\mu(t)$ change over time in-between divisions, and we assume that the growth rate between two consecutive measurements is well approximated by a constant. That is, for a given $\mu_{i,j}$, $\mu(t) \simeq \mu_{i,j}$, for all $t \in [t_{i,j}, t_{i,j+1})$. For every $i$ and some initial value $\nu_i$, using the notation $\delta_{i,j} = t_{i,j+1} - t_{i,j}$ we can now construct the piecewise-linear model

$$\begin{aligned}
\ell(t_{i,1}) &= \nu_i, \\
\ell(t_{i,2}) &= \mu_{i,1}\delta_{i,1} + \ell(t_{i,1}) = \mu_{i,1}\delta_{i,1} + \nu_i, \\
\ell(t_{i,3}) &= \mu_{i,2}\delta_{i,2} + \ell(t_{i,2}) = \mu_{i,2}\delta_{i,2} + \mu_{i,1}\delta_{i,1} + \nu_i, \\
&\vdots \\
\ell(t_{i,n_i}) &= \mu_{i,n_i-1}\delta_{i,n_i-1} + \ell(t_{i,n_i-1}) = \mu_{i,n_i-1}\delta_{i,n_i-1} + \ldots + \mu_{i,1}\delta_{i,1} + \nu_i.
\end{aligned} \tag{S2.12}$$

Note that $\mu_{i,j}$ (that is, the growth rate) is allowed to change from one measurement to the next. Growth rate variations in-between measurements could also be considered (at the price of a more complex inference problem), but this is not justified in our case given the high sampling density (*Materials and methods*).

For all $i$ and $j$, the model unknowns to be inferred from the data $\tilde{\ell}_{i,j}$ are the variable growth rates, $\mu_{i,j}$, and (of less interest here) the cell lengths at the first measurement times after divisions, $\nu_i$. As such, this is an underdetermined problem. To cope with this and with measurement noise, we resort to regularized least squares, a well-established method in the statistical literature [9]. Let us arrange the model parameters into vectors $\underline{\nu} = [\nu_1, \ldots, \nu_m]^T$ and $\underline{\mu} = [\underline{\mu}_1^T, \ldots, \underline{\mu}_m^T]^T$, where $\underline{\mu}_i = [\mu_{i,1}, \ldots, \mu_{i,n_i}]^T$, with $i = 1, \ldots, m$. (For every $i$, entry $\mu_{i,n_i}$, which does not appear in Eq. S2.12, should be understood as the growth rate after the last measurement before cell division). Note in particular that the entries of $\underline{\mu}$ are arranged according to time order. We seek parameters $\underline{\nu}$ and $\underline{\mu} \geq 0$ that minimize

$$(1 - \lambda) \cdot \sum_{i,j} \left(\tilde{\ell}_{i,j} - \Delta_{i,j}^T \cdot \underline{\mu}_i - \nu_i\right)^2 + \lambda \cdot Q(\underline{\mu}), \tag{S2.13}$$

where, for $\Delta_{i,j}^T = [\delta_{i,1}, \ldots, \delta_{i,j-1}, 0, \ldots, 0]$ of size $n_i$, with $j = 1, \ldots, n_i$ and $i = 1, \ldots, m$, $\Delta_{i,j}^T \cdot \underline{\mu}_i + \nu_i$ denotes the predictions of $\ell(t_{i,j})$ obtained from the model of Eq. S2.12 for putative values of $\underline{\nu}$ and $\underline{\mu}$. Term $Q(\underline{\mu}) \geq 0$ denotes a suitable cost function chosen so as to penalize unrealistic fluctuations in the growth rate time-series $\underline{\mu}$, and $\lambda \in (0, 1)$ is a regularization parameter that trades off perfect data fit ($\lambda \simeq 0$) with penalization of fluctuating solutions ($\lambda > 0$). The discussion of the automated choice of $\lambda$ from data is postponed to a dedicated section below. Well-established choices of $Q$ correspond to penalizing the sum of squares of the $d$th (discrete-time) derivative of $\underline{\mu}$ (regarded as a time series), with customary choices $d = 1$ or 2 [9]. Discrete differentiation is expressed by a matrix operation $D_d \cdot \underline{\mu}$, where $D_d$ is an $(n-d) \times n$ matrix. For $d = 2$ (our choice), the $k$th row of $D_2$ has the only nonzero entries 1, $-2$, 1 in positions $k$, $k+1$, $k+2$.

Importantly, for such choice of $Q$, minimization of Eq. S2.13 is a linear least squares problem with bound constraints. For $\underline{\tilde{\ell}} = [\tilde{\ell}_1^T, \ldots, \tilde{\ell}_m^T]^T$, where $\tilde{\ell}_i = [\tilde{\ell}_{i,1}, \ldots, \tilde{\ell}_{i,n_i}]^T$ with $i = 1, \ldots, m$, the estimates sought, denoted by $\hat{\underline{\nu}}$ and $\hat{\underline{\mu}}$, are given by

$$(\hat{\underline{\nu}}, \hat{\underline{\mu}}) = \arg \min_{\underline{\nu}, \underline{\mu} \geq 0} \left\| \begin{bmatrix} \sqrt{(1-\lambda)} \cdot \underline{\tilde{\ell}} \\ 0 \end{bmatrix} - \begin{bmatrix} \sqrt{(1-\lambda)} \cdot \Delta & \sqrt{(1-\lambda)} \cdot L \\ \sqrt{\lambda} \cdot D_2 & 0 \end{bmatrix} \cdot \begin{bmatrix} \underline{\mu} \\ \underline{\nu} \end{bmatrix}, \right\|^2 \qquad \text{(S2.14)}$$

where $\Delta$ is a block-diagonal matrix with blocks defined by $\Delta_i = [\Delta_{i,1}, \ldots, \Delta_{i,n_i}]^T$, $i = 1, \ldots, m$, and $L$ is a block-diagonal matrix with blocks defined by the size-$n_i$ column vectors $L_i = [1, \ldots, 1]^T$, $i = 1, \ldots, m$. Thanks to regularization, the solution of this problem is well-defined, and we solve it via the Python function `lsq_linear` of the `scipy.optimize` module. For a fixed value of $\lambda$ and a given dataset $\underline{\tilde{\ell}}$ with a number of entries in the order of 300 points (some cells being observed over incomplete time periods), optimization takes less than $10^{-2}$ seconds.

**Automatic tuning of regularization parameters** To fix an appropriate value for $\lambda$ directly from the data while ensuring robustness to possible residual segmentation inaccuracies, we used a non-parametric hypothesis testing approach. In conceptual analogy with the well-known discrepancy principle [10], for every mother cell $c$, we choose the largest value of $\lambda$ such that the fitting residuals from the solution of Eq S2.14 do not differ statistically from a representative sample of residuals. This reference set of residuals is calculated under the assumption of balanced growth over a suitable time-interval in the experiment. Assume that, for $i = h, \ldots, k$, mother cell $c$ is in balanced growth in the period $t_{h,1}, \ldots, t_{k,n_k}$. For every value of $i$ in-between $h$ and $k$, we define $e_{i,1}, \ldots, e_{i,n_i}$ as the residuals of a linear fit of the log-length measurements $\tilde{\ell}_{i,1}, \ldots, \tilde{\ell}_{i,n_i}$ at measurement times $t_{i,1}, \ldots, t_{i,n_i}$ (dependence on $c$ is omitted for brevity). Note that these are times in-between cell divisions, where a linear fit for exponential growth is appropriate. The reference residual set is given by

$$\mathscr{R} = \{|e_{i,1}|, \ldots, |e_{i,n_i}| : \; i = h, \ldots, k\}.$$

In practice, we choose the largest $h$ and the smallest $k$ such that, for $i = h, \ldots, k$, the intervals in-between divisions $[t_{i,1}, t_{i,n_i}]$ fall entirely within a regime of balanced growth on glucose. Now define the $\lambda$-dependent fitting residuals from the solution of Eq S2.14, $e_{i,j}^{\lambda} = \tilde{\ell}_{i,j} - \Delta_{i,j}^T \cdot \hat{\underline{\mu}}_i - \hat{\underline{\nu}}_i$, and collect them in the set of residuals at all measurement times

$$\mathscr{R}^{\lambda} = \{|e_{i,1}^{\lambda}|, \ldots, |e_{i,n_i}^{\lambda}| : \; i = 1, \ldots, m\}.$$

We define the estimate $\hat{\lambda}^c$ of $\lambda$ for cell $c$ as the largest value of $\lambda \in (0, 1)$ such that the one-tailed Mann-Whitney $U$ test for $\mathscr{R}^{\lambda}$ larger than $\mathscr{R}$ is negative (*i.e.*, residuals are statistically of similar magnitude) at significance level $\alpha = 0.05$. In Python, this test is implemented by the `mannwhitneyu` function of the `scipy.stats` module. In practice, the search of such $\lambda$ is performed over a sufficiently fine grid in $(0, 1)$. Note that $\hat{\lambda}^c$ is well-defined because (i) for $\lambda = 0$, fitting residuals vanish (overfitting), (ii) for increasing $\lambda$, the magnitude of fitting residuals $e_{i,j}^{\lambda}$

increases, and (iii) for $\lambda$ approaching 1, residuals $\mathscr{R}^{\lambda}$ approach those of a linear fit in-between divisions, which poorly approximates phases of non-balanced growth.

This yields a set of estimates $\hat{\lambda}^c$, one per mother cell $c$. Finally, we establish our choice of $\lambda$ by taking the median of the $\hat{\lambda}^c$ values. Taking the median makes our final choice $\hat{\lambda}$ robust to potential inaccuracies in the definition of the reference residuals set $\mathscr{R}$ and possible specificities of individual mother-cell traces. The execution of the whole procedure over about 100 cells and $n = 300$ data-points (our case) takes around 15 minutes. The regularization parameter $\hat{\lambda}$ determined as explained above was inserted into Eq. S2.14 to estimate growth rate profiles $\hat{\underline{\mu}}^c$ for the various mother cells from the corresponding length sequences $\tilde{\ell}^c$.

**Application to growth medium switches** Regularization in Eq. S2.14 implicitly assumes that no prior information exists on where the most relevant changes in the growth rate time-series $\underline{\mu}$ occur. For cells exposed to growth medium switches, instead, non-smooth growth rate changes are expected to occur after the switching times. This suggests a slight modification of the definition of the regularization term. If the $k$th entry of $\underline{\mu}$ is the growth rate right before a medium switch, we redefine $D_2$ by simply setting to 0 rows $k-1$ and $k$. As a consequence, a non-smooth change in growth rate right after the medium switch is not penalized. Provided this rearrangement of matrix $D_2$ in Eq. S2.14, the calculation of the optimal regularization coefficient $\hat{\lambda}$ first, and of the estimates $\hat{\underline{\mu}}^c$ next are otherwise unchanged.

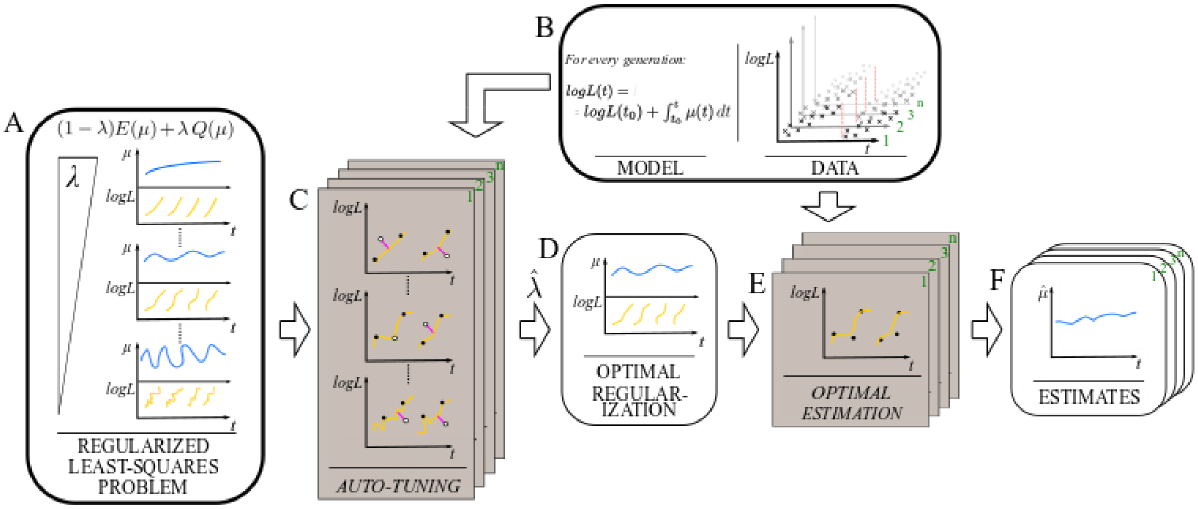


**Figure S2.1: Schematic outline of the method used for the estimation of growth rate from cell length data. A.** Definition of the regularized least-squares problem. $E(\mu)$ represents an error function and $Q(\mu)$ a cost function penalizing large fluctuation in the growth rate, $\lambda$ is the associated regularization parameter. Larger values of $\lambda$ penalize faster fluctuations in the growth rate. **B.** Noisy length measurements in the log domain between divisions are used in a piecewise-linear model to obtain growth rate estimates that may change within a generation. **C-D.** The optimal regularization parameter is chosen for each cell via a hypothesis-testing method and used to obtain the globally optimal regularization parameter over all cells. **E-F.** The optimal regularization parameter along with the defined model is used to fit the data and obtain the sought growth rate estimates for each cell. Rectangular boxes represent procedures. Rounded boxes represent inputs, outputs and results of the method.

#### Estimation of resource allocation profiles from fluorescence data

The description below refers to the estimation problem for a single mother cell $c$. For simplicity, we omit dependence on $c$ from the notation, until this becomes needed. Estimation of the unknown resource allocation profile $\alpha(t)$ relies on the model of Eqs 3-4 in the main text. This is an ODE model for the concentrations of immature and mature reporter proteins ($r_{im}(t)$ and $r_m(t)$) in a single cell.

As explained above, the analysis of single-cell microfluidics data yields noisy measurements of $r_m(t)$ at times $t_{i,j}$. Because the evolution of $r_m$ is assumed to be smooth (no jumps following cell division), *i.e.,* index $i$ plays no special role, we simplify the indexing of data and consider that measurements are taken at (increasing) times $t_k$, with $k = 1, \dots, n$. More precisely, we assume that noisy measurements $y_k$ obey

$$y_k = r_m(t) + \varepsilon_k, \tag{S2.15}$$

where the measurement errors $\varepsilon_k$ form a sequence of independent random variables with mean zero and variance $s_k^2$. We assume that the variance is estimated from the data (*Materials and methods*).

Let $x(t) = [r_{im}(t), r_m(t)]^T$. The model of Eqs 3-4 can be written in the form

$$\dot{x}(t) = F(t)x(t) + G(t)\alpha(t), \tag{S2.16}$$

where $\alpha(t)$ is the unknown profile to be inferred from $y_1, \dots, y_n$. Matrices $F(t)$ and $G(t)$ are given in Eq 9. They depend on parameters $\gamma$ and $k_{mat}$ whose values are known (Tab. S3), $\beta$, which only appears as a rescaling of $\alpha(t)$, and the growth rate profile $\mu(t)$. The parameter $\beta$ cannot be determined from the data, so we drop it from the model and bear in mind that estimation of $\alpha(t)$ below is in fact estimation of $\alpha(t)/\beta$. For $\mu(t)$ known, Eq. S2.16 is a linear state-space model. To leverage this property, we pretend that $\mu(t)$ is known. In practice $\mu(t)$ will be replaced by estimates $\hat{\mu}(t)$ obtained by interpolation of the vector estimate $\underline{\hat{\mu}}$ from the previous section.

Reconstruction of $\alpha(t)$ from sampled, noisy measurements $y_1, ..., y_n$ is an ill-posed problem. To obtain relevant, robust estimates we resort again to regularization. Here, the standard, linear state-space form of model S2.15–S2.16 prompts us to pose estimation as a Bayesian problem, and solve it with effective tools based on Kalman filtering [11]. Similar to [7], we introduce a probabilistic prior on $\alpha$ in the form of the Ornstein-Uhlenbeck process

$$d\alpha(t) = -\theta \cdot \alpha(t)dt + \sigma \cdot dW(t), \tag{S2.17}$$

where $W(t)$ is a standard Wiener process. We additionally assume that $\alpha(0)$ has a zero-mean Gaussian distribution with variance $\sigma^2/(2\theta)$, which implies that the process has stationary statistics. In simple words, this prior introduces information about the profile sought by assigning different probabilities to different profiles, depending on parameters $\theta$ and $\sigma$. Large (small) values of the ratio $\sigma/\sqrt{\theta}$ favor larger (small) fluctuations of $\alpha$. Large (resp. small) values of $\theta$ favor fast (slow) fluctuations. This information is quantitatively conveyed by the autocovariance function of the process [12], which has the form

$$\mathbb{E}[\alpha(t)\alpha(\tau)] = \sigma^2/(2\theta) \cdot \exp(-\theta \cdot |t - \tau|), \tag{S2.18}$$

for any two times $t, \tau$, where the first factor is the process variance and the second factor shows the exponentially decaying "memory" of the process. We postpone the illustration of the automated choice of $\theta$ and $\sigma$, which play the role of regularization parameters, to the next section.

Given the probabilistic prior, the problem of calculating an optimal estimate $\hat{\alpha}(t)$ for $\alpha(t)$ given data $y_1, \ldots, y_n$ becomes the Bayesian problem of calculating the conditional expectation

$$\hat{\alpha}(t) = \mathbb{E}[\alpha(t)|y_1, \ldots, y_n]. \tag{S2.19}$$

This reconstruction is optimal in the sense that it minimizes the variance of the estimation error at any time $t$ [12]. To compute the right-hand side of Eq. S2.19, we restrict attention to estimates $\hat{\alpha}(t)$ at times $t_k$, and leverage linearity properties of Eqs S2.15–S2.17 to recast the problem into the form of an efficient computation known as Kalman smoothing [13]. Define the augmented state $\xi(t) = [x(t)^T \ \alpha(t)]^T$. From the previous relations one can write the linear stochastic state-space model

$$\begin{aligned} d\xi(t) &= \begin{bmatrix} F(t) & G(t) \\ 0 & -\theta \end{bmatrix} \xi(t)dt + \begin{bmatrix} 0 \\ \sigma \end{bmatrix} dW(t), \\ y_k &= C\xi(t_k) + \varepsilon_k, \end{aligned} \tag{S2.20}$$

where the zeros denote null matrix blocks of appropriate dimensions, and $C = [0, 1, 0]$. For generic indices define

$$\hat{\xi}_{k|h} = \mathbb{E}[\xi(t_k)|y_1, \ldots, y_h], \qquad P_{k|h} = \mathbb{E}[(\xi(t_k) - \hat{\xi}_{k|h})(\xi(t_k) - \hat{\xi}_{k|h})^T],$$

where $\hat{\xi}_{1|0}$ and $P_{1|0}$ are in particular the prior mean and covariance matrix of $\xi(t_1)$.

The Kalman filter is a forward iteration to calculate $\hat{\xi}_{k|k}$ (and $\hat{\xi}_{k|k-1}$) along with $P_{k|k}$ (and $P_{k|k-1}$) for $k$ from 1 to $n$. On top of this, the Kalman smoother is obtained by further running a backward iteration that calculates $\hat{\xi}_{k|n}$ along with $P_{k|n}$ for $k$ from $n$ down to 1. This provides estimates of all entries of $\xi$. In particular, estimate $\hat{\alpha}(t)$ at time $t_k$ and the corresponding estimation error variance is then obtained by the equations

$$\hat{\alpha}(t_k) = S\hat{\xi}_{k|n}, \qquad \mathbb{E}[(\hat{\alpha}(t_k) - \alpha(t_k))^2] = SP_{k|n}S^T, \tag{S2.21}$$

with $S = [0, 0, 1]$. As the terminology suggests, $\hat{\alpha}$ will display smooth dynamics, which agrees with expectations except at times where the culture medium is switched. The adaptations of the approach to cope with this are described below.

The Kalman filtering equations are standard (see details in [13, 14]). For every $k$, two steps are performed. The prediction step consists in calculating $\hat{\xi}_{k+1|k}$ and $P_{k+1|k}$ from $\hat{\xi}_{k|k}$ and $P_{k|k}$ by the solution of the system dynamics given by S2.20 and a corresponding Riccati equation from time $t_k$ to $t_{k+1}$. The update step consists in integrating the information from the new measurement $y_{k+1}$ to get $\hat{\xi}_{k+1|k+1}$ and $P_{k+1|k+1}$ from $\hat{\xi}_{k+1|k}$ and $P_{k+1|k}$ via simple algebraic operations involving Eq. S2.15. This iteration is performed for $k = 1, \ldots, n-1$, starting from $\hat{\xi}_{1|0}$ and $P_{1|0}$ (at $k = 1$, the prediction step is skipped). The less known smoothing iteration is also standard, and consists purely in algebraic calculations on data and quantities calculated in the forward filtering iteration. The choice of initial conditions $\hat{\xi}_{1|0}$ and $P_{1|0}$ (the prior on $\xi(t_1)$)

is determined in part by the prior on $\alpha$. We set $\hat{\xi}_{1|0} = [0,0,0]^T$ and $P_{1|0} = \mathrm{diag}\big(M, M, \sigma^2/(2\theta)\big)$, with $M$ large ($10^5$), corresponding to virtual lack of information on $r_{im}$ and $r_m$ prior to the experiments.

The computational complexity of the whole procedure is linear in the number of data points $n$, and is mostly determined by the inversion of small ($3 \times 3$) matrices. For a single mother cell data series with a number of points $n$ in the order of 300 (our case), our Python implementation using function `odeint` of the `scipy.integrate` module for numerical integration takes about 1 second. If the Gaussian assumptions are violated, the estimates computed by this procedure have the interpretation of optimal estimates (minimal error variance) in the class of linear functions of the data [13]. Thanks to this, the method is robust to moderate deviations from Gaussianity. Computational efficiency of the method is especially important for the automated tuning of parameters $\theta$ and $\sigma$, as explained below. A demonstration of the effectiveness of the approach was given in a validation study with synthetic data (*Materials and methods* and Fig. S18).

**Automatic tuning of regularization parameters** To fix the prior parameters we follow a maximum-likelihood approach. That is, we seek parameters $\theta$ and $\sigma$ that maximize

$$f_{\theta,\sigma}(y_1,\ldots,y_n) = f_{\theta,\sigma}(y_1) \cdot \prod_{k=2}^{n} f_{\theta,\sigma}(y_k|y_1,\ldots,y_{k-1}),$$

where the notation indicates probability densities under candidate values of $\theta$ and $\sigma$, and the factorization of the joint density of the data is a consequence of Bayes' law. Maximization of the likelihood is equivalent to minimization of the negative log-likelihood. Under Gaussianity of the prior and of measurement noise, from the equation above and Eq. S2.15, the negative log-likelihood is, up to constant additive terms, given by

$$\mathscr{L}(\theta,\sigma|y_1,\ldots,y_n) = \sum_{k=1}^{n} \log\Lambda_k(\theta,\sigma) + \frac{1}{2}\left[\frac{y_k - \hat{y}_{k|k-1}(\theta,\sigma)}{\Lambda_k(\theta,\sigma)}\right]^2,$$

where $\hat{y}_{k|k-1}(\theta,\sigma)$ and $\Lambda_k(\theta,\sigma)$ are, respectively, the mean and variance of the conditional distribution $f_{\theta,\sigma}(y_k|y_1,\ldots,y_{k-1})$, with $k = 2,\ldots,n$, while $\hat{y}_{1|0}(\theta,\sigma)$ and $\Lambda_1(\theta,\sigma)$ are those of $f_{\theta,\sigma}(y_1)$. It follows from the Kalman filtering theory that $\hat{y}_{k|k-1}(\theta,\sigma) = C\hat{\xi}_{k|k-1}(\theta,\sigma)$ and $\Lambda_k(\theta,\sigma) = CP_{k|k-1}(\theta,\sigma)C^T + s_k^2$, with $k = 1,\ldots,n$, where $\hat{\xi}_{k|k-1}(\theta,\sigma)$ and $P_{k|k-1}(\theta,\sigma)$ are the one-step state predictions and covariance matrices obtained in the Kalman filtering iteration described above, run for the given values of $\theta$ and $\sigma$ [13].

By the machinery above, for every mother cell $c$, we obtain estimates of $\theta$ and $\sigma$ by the optimization

$$(\hat{\theta}^c, \hat{\sigma}^c) = \arg\min_{\theta>0,\sigma>0} \mathscr{L}(\theta,\sigma|y_1^c,\ldots,y_n^c),$$

where $y_1^c,\ldots,y_n^c$ is the fluorescence time-series for mother cell $c$. In practice, optimization is solved numerically by the Python function `minimize` of the `scipy.optimize` module. In this optimization, objective function $\mathscr{L}(\theta,\sigma|y_1^c,\ldots,y_n^c)$ is evaluated at every explored value of $\theta$ and $\sigma$ by running the corresponding Kalman filter iteration on data $y_1^c,\ldots,y_n^c$. In total, in a typical experiment with 100 cells and for $n = 300$, the computation takes around 2 hours.

Finally, our choice for parameters $\theta$ and $\sigma$ is defined by the median of the values $\hat{\theta}^c$ and $\hat{\sigma}^c$ obtained over all mother cells $c$. This single choice, $\hat{\theta}$ and $\hat{\sigma}$, is plugged into the Kalman filter, which is then applied separately to all cells $c$ in an experiment to get estimates $\hat{\alpha}^c$ of the resource allocation profile from the corresponding fluorescence time-series $y_1^c, \ldots, y_n^c$.

**Application to growth medium switches** For the estimation of resource allocation strategies over switches of the culture medium, considerations similar to those for the estimation of growth rate apply. At switching times, resource allocation may change over a short time interval. Smooth reconstruction of $\alpha$ at these times may therefore be artifactual. In essence, this is a consequence of the stationarity of the prior autocovariance function Eq. S2.18, which stipulates that the dependence between a "current" value $\alpha(t)$ and its "future" evolution $\alpha(\tau)$ $(\tau > t)$ is only a function of the time-lag $\tau - t$, irrespective of the specific value of $t$. Whereas this uniform behavior is relevant for a given medium, the relation between $\alpha(t)$ and $\alpha(\tau)$ is expected to be looser across a medium switch due to changed conditions.

To cope with this, rather than introducing arbitrary (and technically cumbersome) local changes in the prior on $\alpha$, we separate out the reconstruction problem over different growth conditions. The procedure outlined hereafter applies equally to all mother cells $c$ in the analysis, both in the calculation of optimal regularization parameters and in the subsequent calculation of optimal estimates of $\alpha(t)$.

Let $q_h$, with $h = 1, \ldots, M-1$, be the (increasing sequence of) indices such that $t_{q_h}$ are the measurement times right before (or equal to) the time of a medium switch. $M$ is the number of medium switches over the whole experimental horizon. Note that these times are unrelated with cell division times. Also let $q_0 = 0$ and $q_M = n$. Then, for the relevant values $\sigma$ and $\theta$:

- For $h = 0, \ldots, M-1$: define $\hat{\xi}_{q_h+1|q_h} \triangleq [0,0,0]^T$ and $P_{q_h+1|q_h} \triangleq \mathrm{diag}\big(M, M, \sigma^2/(2\theta)\big)$ as the prior on $\xi(t_{q_h+1})$, and run the Kalman filering-smoothing procedure over indices $(q_h+1, \ldots, q_{h+1})$. This yields estimates $\hat{\xi}_{q_h+1|q_{h+1}}, \hat{\xi}_{q_h+2|q_{h+1}}, \ldots, \hat{\xi}_{q_{h+1}|q_{h+1}}$ and estimation error covariance matrices $P_{q_h+1|q_h+1}, P_{q_h+2|q_h+1}, \ldots, P_{q_{h+1}|q_{h+1}}$ computed solely from the measurements $y_{q_h+1}, \ldots, y_{q_{h+1}}$.

- Return the whole sequence of estimates

  $$(\hat{\xi}_{1|q_1}, \ldots, \hat{\xi}_{q_1|q_1}), \ldots, (\hat{\xi}_{q_h+1|q_{h+1}}, \ldots, \hat{\xi}_{q_{h+1}|q_{h+1}}), \ldots, (\hat{\xi}_{q_{M-1}+1|q_M}, \ldots, \hat{\xi}_{q_M|q_M})$$

  and corresponding estimation error covariance matrices

  $$(P_{1|q_1}, \ldots, P_{q_1|q_1}), \ldots, (P_{q_h+1|q_{h+1}}, \ldots, P_{q_{h+1}|q_{h+1}}), \ldots, (P_{q_{M-1}+1|q_M}, \ldots, P_{q_M|q_M}).$$

The fact that, in every period $h$, only measurements from the same period are used breaks the dependence on the dynamics in nearby periods and enables sudden changes at medium switch times.

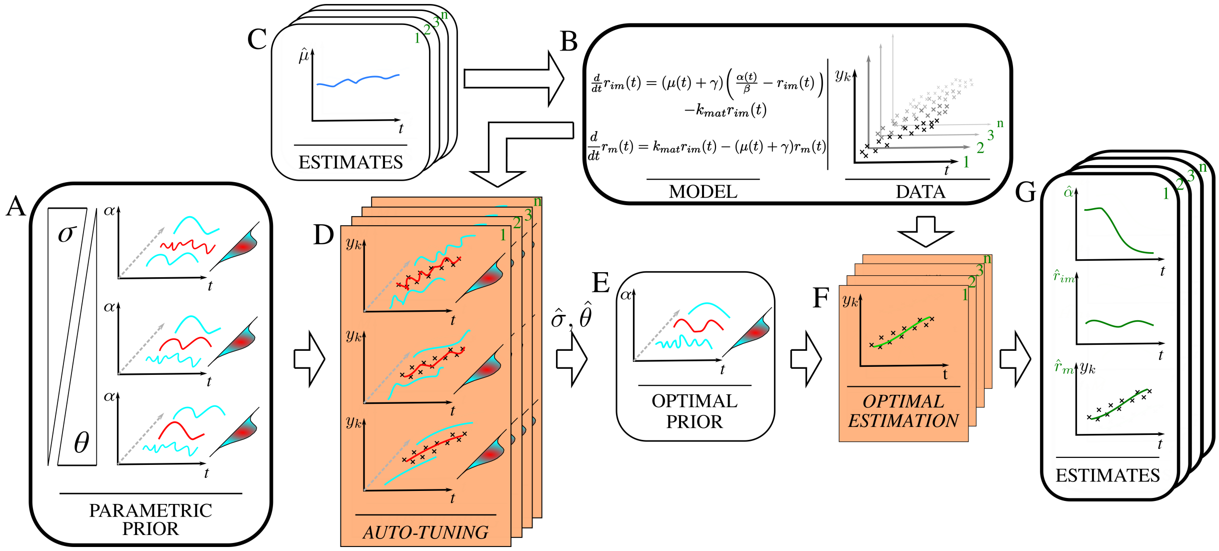


**Figure S2.2: Schematic outline of the Bayesian approach for estimating single-cell resource allocation profiles. A.** Probabilistic priors on $\alpha$. A larger $\sigma$ (smaller $\theta$) assigns higher probability to faster fluctuations in the profile. **B-D.** The kinetic model, the fluorescence data of individual cells, along with their corresponding estimated growth rate (Fig. S2.1) are used in the auto-tuning step so that the best values of $\sigma$ and $\theta$ are selected by comparing the fluorescence profiles predicted by the model with the data via a maximum-likelihood approach. **E-G.** The optimal prior selected is used with the single-cell fluorescence intensity measurements for the optimal estimation of individual resource allocation profiles and predicted fluorescence outputs. Rectangular boxes represent procedures. Rounded boxes represent inputs, outputs and results of the method.

### Text S3 Model analysis

The aim of this supplementary text is to demonstrate a consequence of the ribosomal resource allocation model that can be confronted with the experimental data: if resource allocation ($\alpha$ or $\alpha/\beta$) is higher during the upshift, then the ribosome concentration $r$ will also be higher in the same time-interval.

We remind the model of ribosomal resource allocation (Eq. 1 in the main text):

$$\frac{d}{dt}r(t) = (\mu(t) + \gamma) \left( \frac{\alpha(t)}{\beta} - r(t) \right). \tag{S3.22}$$

The growth rate can be expressed in terms of the total protein synthesis rate (Eq. S1.5 in Text S1):

$$\mu(t) + \gamma = \beta \, v_{prot}(t). \tag{S3.23}$$

Moreover, with the definition of the protein synthesis rate in the main text as $v_{prot}(t) = r_{act}(t)\,\sigma$, the product of the concentration of active ribosomes and their activity, where $r(t) = r_{act}(t) + r_{inact}(t)$, we obtain (Eq. 2 in the main text):

$$\mu(t) + \gamma = \frac{r_{act}(t)}{\beta}\,\sigma. \tag{S3.24}$$

Assuming that a fraction $\delta$ of the total ribosome pool is active, this equation becomes

$$\mu(t) + \gamma = \frac{\delta\, r(t)}{\beta}\,\sigma. \tag{S3.25}$$

Combining all of the above equations, we find

$$\frac{d}{dt}r(t) = (\alpha(t) - \beta\, r(t))\,\delta\, r(t)\,\sigma. \tag{S3.26}$$

For constant $\alpha(t) = \alpha^*$, the above equation reduces to a logistic equation [15]. This model has the following explicit solution for $t \geq 0$, where $r(0) = r_0$ is the ribosome concentration just before the upshift:

$$r(t) = \frac{\alpha^*/\beta}{1 + \left( \frac{\alpha^*/\beta}{r_0} - 1 \right) e^{-\alpha^*\,\sigma\,\delta\,t}}. \tag{S3.27}$$

As expected, when $t \to \infty$, the solution of $r$ approaches $\alpha^*/\beta$. Let $r^+(t)$ be the solution corresponding to the case of $\alpha^{*,+} > \alpha^*$, where $r_0^+ \geq r_0$. It can be shown by means of the above equation that $r^+(t) > r(t)$, for $t \in [0, T]$ and $T > 0$. That is, a higher allocation of resources of ribosomes during the upshift leads to a higher ribosome concentration over the whole duration of the upshift.

The data in Fig. 4 in the main text indicate that $\alpha$ is not necessarily constant after an upshift. In this case, the model of Eq. S3.28 does not have an explicit solution. A similar result as above, however, can be obtained from basic results in dynamical systems theory [16]. Let $\alpha^+(t) > \alpha(t)$, for $t \in [0, T]$. Moreover, denote the right-hand side of Eq. S3.28 by $f_r(r(t), \alpha(t))$. We now define a so-called super-solution $r^+(t)$:

$$\frac{d}{dt} r^+(t) = f_r(r^+(t), \alpha^+(t)), \tag{S3.28}$$

where $f_r(r^+(t), \alpha^+(t)) > f_r(r^+(t), \alpha(t))$, for $t \in [0, T]$. From Lemma 1.2 in [16], whenever $r_0^+ > r_0$, it follows that $r^+(t) > r(t)$. In other words, in the case of time-varying $\alpha$, a higher allocation of resources of ribosomes during the upshift leads to a higher ribosome concentration over the whole duration of the upshift as well. These observations agree with the experimental data shown in Fig. S21, as discussed in the main text.
